# Niche constraints drive differences between mycorrhizal fungal guilds in future range shifts

**DOI:** 10.64898/2026.05.18.725971

**Authors:** Joseph D. Edwards, Johan van den Hoogen, Christine V. Hawkes, Mark A. Anthony, Camille S. Delavaux, Justin D. Stewart, Kathleen K. Treseder, Xiao Feng, Monica Papeş, Robert Muscarella, Petr Kohout, Petr Baldrian, E. Toby Kiers, Thomas C. Lauber, P. Brandon Matheny, Thomas W. Crowther, Michael E. Van Nuland, Clara Qin, Colin Averill, Stephanie N. Kivlin

## Abstract

Mycorrhizal fungi are a diverse and ubiquitous group of plant symbionts whose distribution strongly influences ecosystem function across the globe. Yet, until now, we do not have quantitative data on the range sizes of different mycorrhizal fungal taxa, limiting our capacity to forecast future shifts in community composition and function. Here, we use 115,924 DNA sequence-derived observations to map the distribution of 651 common mycorrhizal fungal taxa and forecast future changes to their range sizes. We demonstrate that climate, plant cover, and soil factors, particularly mean annual temperature, net primary productivity, and soil organic carbon, exert major control over mycorrhizal fungal distributions. Based on these drivers, the ecological niches of mycorrhizal fungi consistently differ between arbuscular and ectomycorrhizal functional guilds. Arbuscular mycorrhizal fungal taxa generally occupy a wider niche breadth than ectomycorrhizal fungi, occurring across larger ranges of climate, soil, plant cover, topography, and disturbance conditions. Our models also predict widespread decreases in the suitable range size of mycorrhizal fungal taxa under projected future global climates, with average ranges decreasing by 13.8% or ∼2.2 million km^2^ under high emissions scenarios (ssp5-8.5). This decrease in projected range size will be most pronounced for ectomycorrhizal fungi, strongly linked to constraints from their smaller overall niches. By generating a global atlas of common mycorrhizal fungi and their associated environmental niche, we establish a critical baseline for widely suspected declines in global fungal biodiversity.

**Significance Statement:** Organismal niches are defined by the environments they inhabit and can shape how they respond to future changes. Here, we characterize the realized niche space for over 600 taxa of mycorrhizal fungi, a critical group of ecosystem engineers, and use these traits to predict their responses to future climate scenarios. We found that the strength of environmental drivers in predicting taxon occurrence strongly differs between mycorrhizal guilds, with these differences explaining their divergent projected range shifts. These findings show that mycorrhizal fungal distributions are structured by specific biotic and abiotic cues, with the potential to inform how changes to mycorrhizal fungal biodiversity are anticipated and managed over the coming century.

## Introduction

Understanding the environmental conditions influencing an organism’s realized niche is critical to forecasting the effects of global change on species distributions. Individual species and functional groups vary in niche breadth and sensitivity to perturbations (1). However, most niche characterizations focus on aboveground organisms due to the challenges of considering belowground obligate symbiotes (2–5). One key group of belowground organisms are mycorrhizal fungi. These fungi are abundant and widespread terrestrial symbionts, occurring on all continents and in 90% of plant species (6). Mycorrhizal fungi play an important role in regulating aboveground biodiversity and Earth’s ecological health (7) with the potential to receive 6 - 13% of total carbon (C) fixed by plants (8) and facilitating up to 80% of otherwise plant-limiting nutrient uptake to their hosts (9). Given their ecological importance, delineating the drivers of mycorrhizal fungal niches across species and functional guilds is a crucial goal for biodiversity conservation (10).

Globally, two mycorrhizal guilds account for the majority of mycorrhizal associations: arbuscular mycorrhizal (AM) fungi and ectomycorrhizal (EcM) fungi (11). AM fungi are an ancient, monophyletic lineage that is thought to be relatively species poor with ∼1000 estimated species globally (12). EcM fungi are a diverse group that evolved independently more than 80 times from saprotrophic ancestors (13, 14), with some projections suggesting up to 25,000 - 55,000 species (15–17). In addition, some EcM fungi possess a metabolic toolbox for decaying organic matter (18–20). These differences between AM and EcM fungi drive ecosystem-level “syndromes” where dominant mycorrhizal guilds can alter multiple above- and belowground functions (21–25) and ecosystem responses to global change factors (26–32).

There is limited evidence directly comparing the realized niche dimensions (*i.e.,* observed environment) between AM and EcM fungal taxa. While AM fungal geographic range sizes are consistently larger than for EcM fungi (33), mapping range sizes alone is not sufficient to understand underlying environmental niche determinants. Niche realization in obligate symbionts is dependent on the balance between biotic (host-driven) and abiotic (*e.g.,* climate, soil) influences. Host specificity is generally greater for EcM than AM fungi (34). Consequently, the higher prevalence of host-generalist AM fungi indicates that their distributions are less tightly constrained by host identity, and more strongly influenced by abiotic conditions or spatial processes. In contrast, the narrower host associations in EcM fungi can impose stronger biotic filtering on their realized niches (35, 36). Climate factors also help determine AM and EcM fungal niches, with strong sensitivity to temperature and precipitation across both guilds (37–40). However, community-level analysis reveals that the relative influence of specific climatic factors on mycorrhizal fungal niche breadth varies across latitudes, with mycorrhizal fungal taxa occupying broader temperature niches but narrower precipitation niches at higher latitudes (41). In addition to climate, soil factors like pH, organic carbon, and nutrients often associate with mycorrhizal fungal distributions (42–45) and intra-guild variation in optimal edaphic conditions can be high for both AM and EcM fungi (46, 47). Despite the clear influence of biotic and abiotic environmental conditions for determining mycorrhizal fungal habitat suitability, the relative strength of these factors across and within AM and EcM fungal guilds remains unknown.

Assessing how mycorrhizal fungal realized niches will respond to global change is essential to developing specific goals for safeguarding above and belowground biodiversity. Our understanding of mycorrhizal fungal realized niches largely relies on observations of their geographic distributions (48) that are directly influenced by climate at large spatial scales (49). Thus, changes to temperature and precipitation regimes over the next century are likely to severely impact their geographic distributions (27, 50–52). However, an organism’s current realized niche breadth and determinants may provide key insights into future realized niche space (53). Forecasting temporal shifts in species occurrences by forward projecting their niche models with expected future climate data often predicts significant changes to species distribution ranges (54–57). Critically, niche determinants of plant-associated fungi can fundamentally differ from their hosts (58), increasing risk of future mismatches in mutualist responses to changing environments (59).

Here, we aim to determine environmental predictors of mycorrhizal fungal distributions within and between mycorrhizal fungal guilds. We used an extensive database of 115,924 georeferenced samples of mycorrhizal fungal genetic taxon occurrences (Figure 1; 60, 61, 62) to map the geographic distributions and ecological niches of 249 AM fungal virtual taxa (VTX) and 402 EcM fungal species. We then project these distributions (*i.e.,* applied fitted models to spatial gridded environmental layers) into a worldwide atlas to predict areas of high occurrence probability for these mycorrhizal fungal taxa (Figure S1). Using these projections, we also quantify how mycorrhizal fungal range suitability may respond to “best” (ssp1-2.6) and “worst” (ssp5-8.5) case future global change scenarios. We used these global mycorrhizal fungal taxa occurrence databases and machine-learning niche models to test three major hypotheses focused on the differences between AM and EcM fungal guilds in their: (1) realized niche breadth, (2) strength of environmental correlates in their distribution, and (3) potential sensitivity of their distribution to future climate conditions. Specifically, we hypothesized that (H1) Mycorrhizal fungal guilds occupy distinct niches, with broader realized niches for AM fungi than EcM fungi based on their wider geographic distributions and more numerous plant species association; (H2) The relative importance of environmental features in explaining occurrence probability differs between AM and EcM fungal taxa, with wider environmental tolerances for AM fungi leading to greater importance for other variables (soil properties, vegetation/ land cover, physical properties, and disturbance) while narrower tolerance for EcM fungi will lead to stronger association with abiotic variables (climate/ soil); and, (H3) Mycorrhizal fungal taxa with narrower niches will be more sensitive to future climate conditions, with greater proportional changes in their predicted range sizes under altered climate conditions. Testing these hypotheses will help fill critical gaps in our basic understanding of the fundamental differences in the environmental drivers for mycorrhizal fungal distributions and their potential vulnerability to global change.

**Figure 1.**
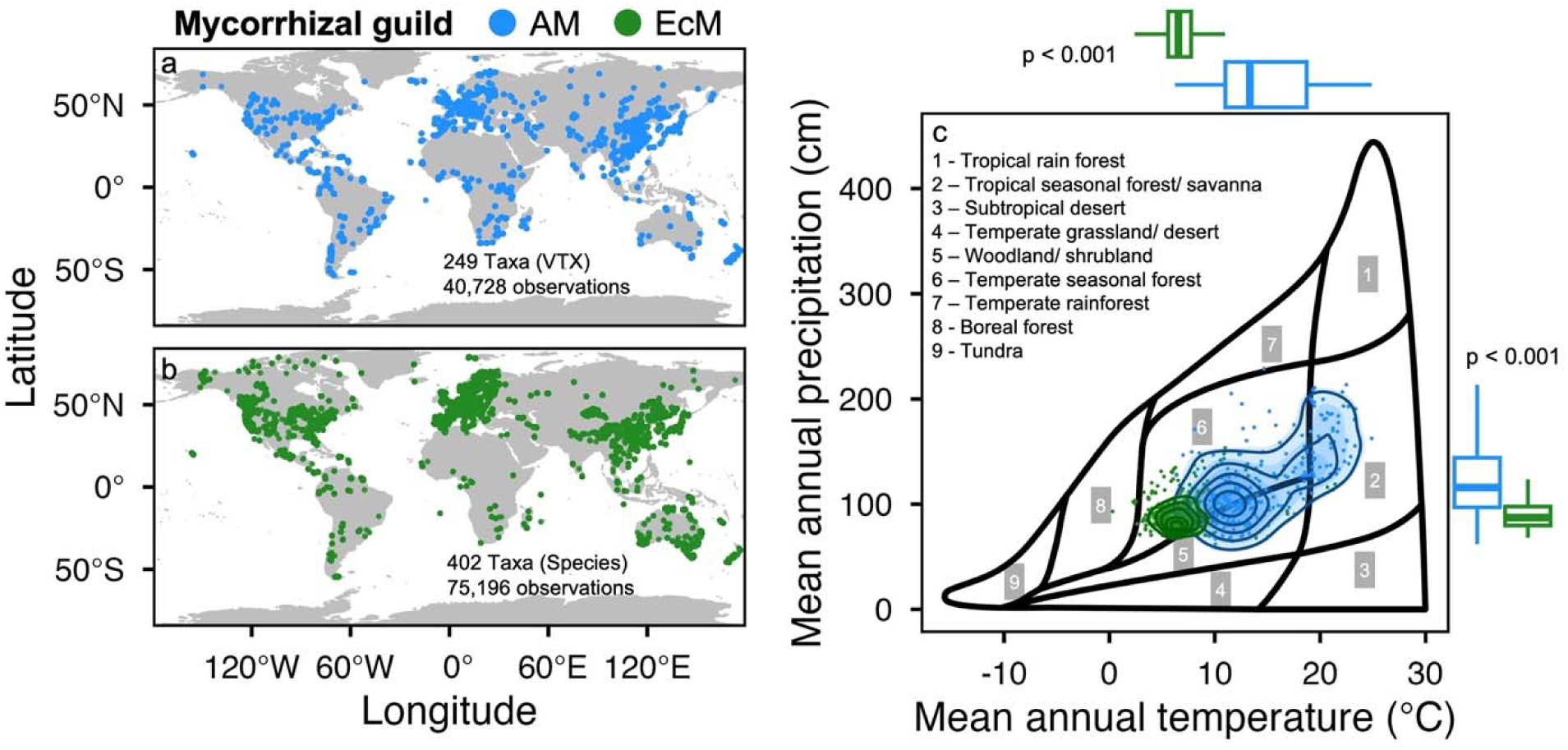
Map of georeferenced sample locations for arbuscular mycorrhizal AM (a; blue) and ectomycorrhizal EcM (b; green) fungal taxa. Mean climatic niches of AM and EcM fungi overlaid on Whittaker’s biome distributions (c). Each point represents a fungal taxon’s mean environmental condition at a sampled location. AM fungi tend to occur in warmer and wetter biomes such as tropical seasonal forests and savannas (2), while EcM fungi are centered around colder temperate seasonal forests and boreal forests (6 & 8).

## Results

### H1 Realized niche breadth

In support of our first hypothesis that mycorrhizal fungi inhabit distinct realized niches, we found AM and EcM fungal taxa generally differed in the average climate in which they were observed (Figure 1). Based on occurrences within our sampled regions, AM fungal taxa (VTX) frequently have mean climate niches in warmer (median = 13.3 °C MAT, IQR = 7.7 °C) and wetter (median = 115.8 cm MAP, IQR = 47.1 cm) biomes, such as temperate seasonal forests and tropical seasonal forests and savannas (Figure 1). By contrast, mean climate niches of EcM fungal taxa (species) occurred predominantly in colder (median = 6.5 °C MAT, IQR = 2.2 °C) and drier (median = 87.2 cm MAP, IQR = 18.0 cm) biomes, such as boreal forests and cold temperate seasonal forests.

In support of our hypothesis that AM fungi inhabited broader realized niches than EcM fungi, the diversity of environmental conditions occupied by mycorrhizal fungal taxa differs between mycorrhizal guilds (Figure 2). When we compared the total range of environmental conditions (*e.g.,* maximum MAT versus minimum MAT) occupied by each taxon, we found that, on average, AM fungal taxa were distributed across a wider range of climatic variables than EcM fungal taxa (Wilcoxon rank-sum r = 0.26 – 0.78, p < 0.001). For example, the range of MAT in AM fungal taxa occurrences was on average 15.3% greater than for EcM fungal taxa (r = 0.18, p < 0.001). The AM taxon with the smallest observed annual temperature range (*VTX00268*, *Glomus sp.*) spanned a breadth of 17.7 °C with a minimum (min) of 9.8 °C and a maximum (max) of 27.6 °C. The range is almost three times as much as the EcM fungal taxon with the smallest observed MAT range (*Inocybe semifulva*, range: 6.4 °C, min: 4.7 °C, max: 11.0 °C).

**Figure 2.**
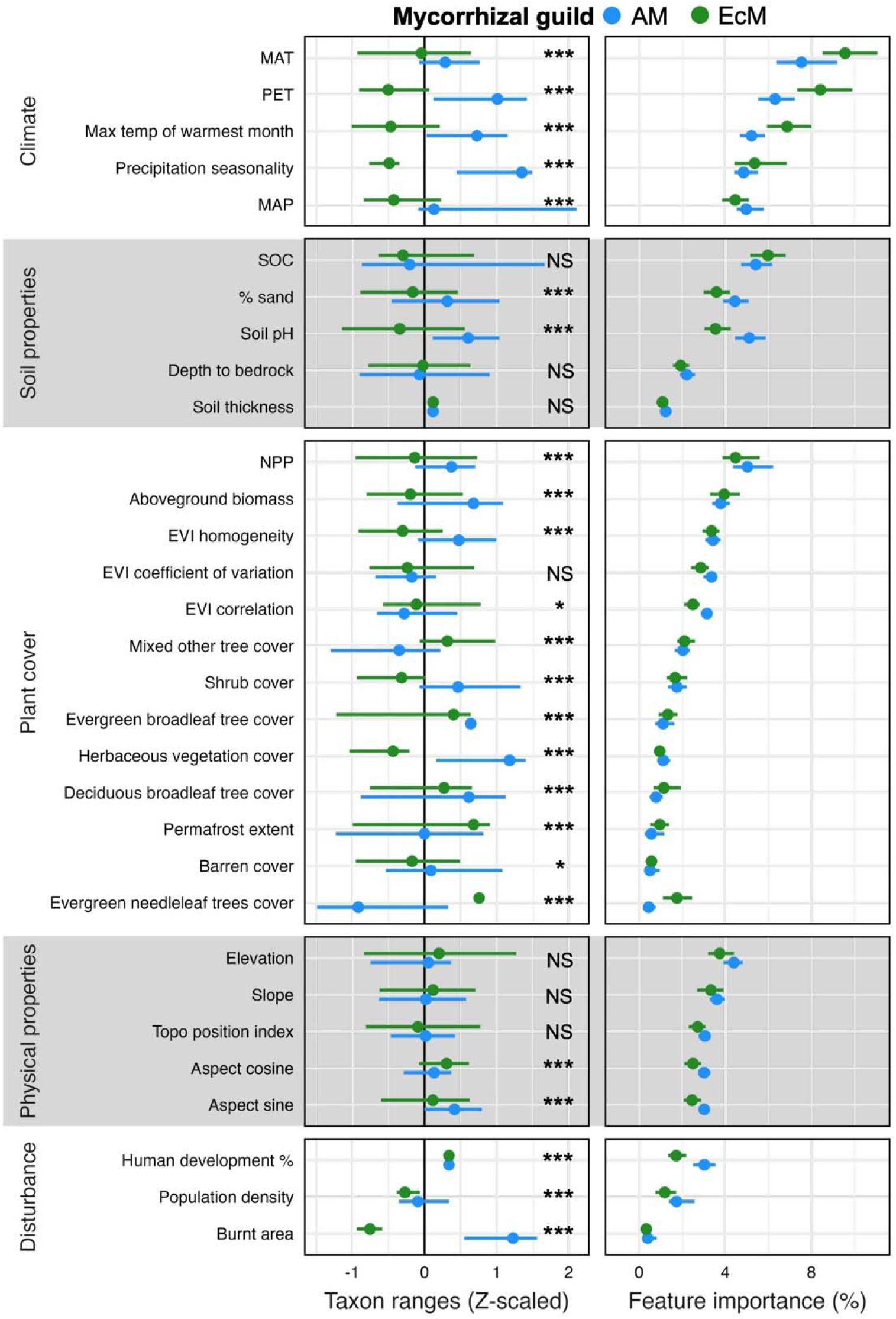
Ecological niche axes for arbuscular mycorrhizal (AM; blue) and ectomycorrhizal (EcM; green) fungal taxa separated into climatic, disturbance, physical properties, plant cover, and soil derived from environmental covariates for each sampled location. The median range of variable values (maximum – minimum) for each taxon and inter-quartile ranges among taxa (z-transformed) are displayed on the left and the median relative feature importance (%) of these variables derived from global random forest niche models for each mycorrhizal fungal taxon is shown on the right. Models were constructed using the Google Earth Engine random forest classifier and variable importance was assessed as mean decrease in node impurity (Gini importance). Significant differences between mycorrhizal guilds for variable ranges are indicated as p>0.05 (NS), p<0.05(*), p<0.01(**), and p<0.001(***) based on Wilcoxon rank-sum test to account for non-normal distribution of variable values and uneven sample size for AM (n = 249 taxa) and EcM (n = 402 taxa) fungi.

**Figure 3.**
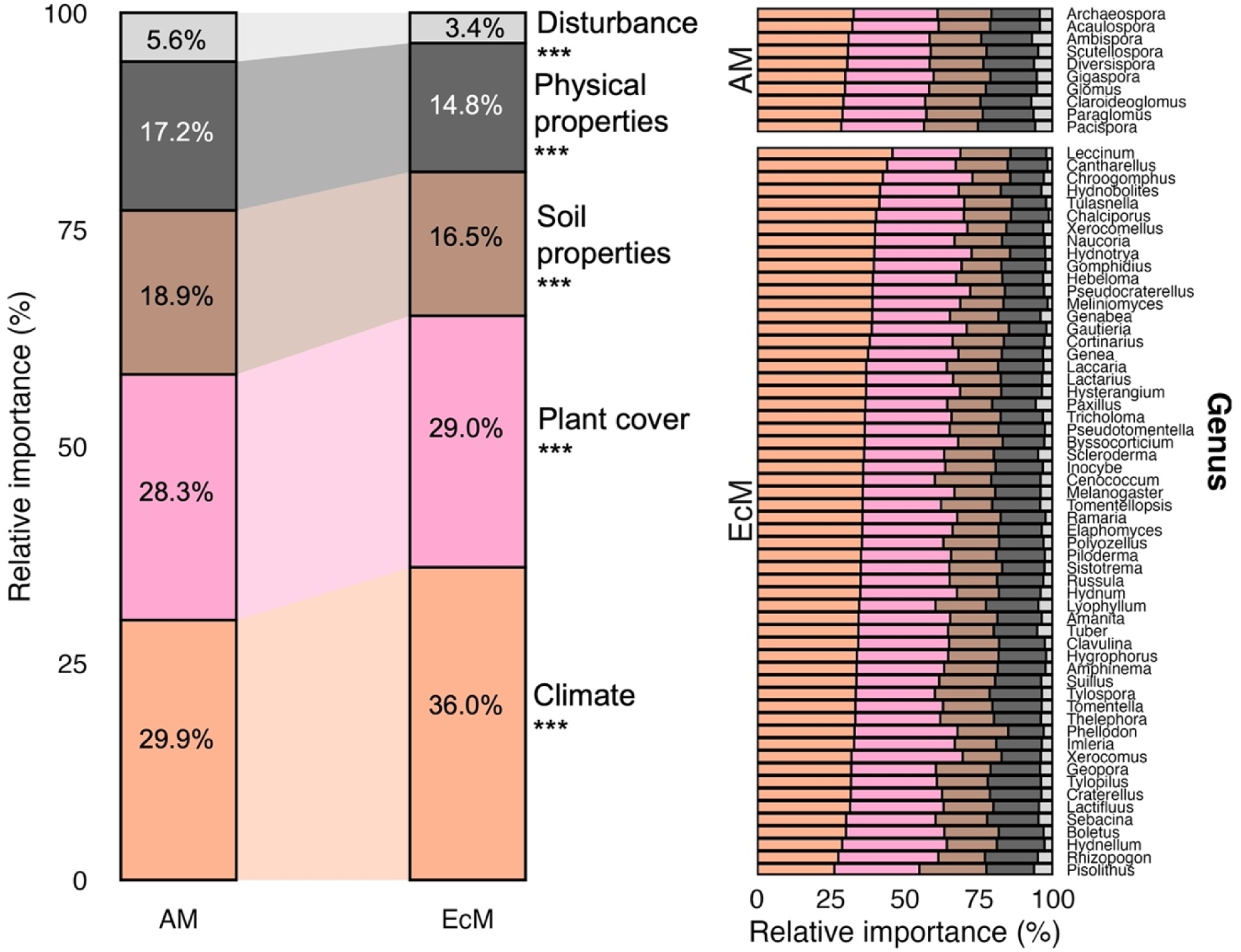
Average relative feature importance of climate (orange), plant cover (pink), soil properties (brown), physical properties (black), and disturbance (grey) of all AM fungi and all EcM fungi shown on the left side and genus-level resolution within each guild shown on the right. Significant differences in the relative variation explained by these features between mycorrhizal guilds are indicated as *** (p < 0.001) based on a Wilcoxon rank-sum test. AM fungi generally have less of their niche explained by climatic variables and plant cover compared to EcM fungi. AM fungi also have more of their niche ascribed to soil properties, physical properties, and disturbance factors than EcM fungi. Across genera, EcM fungi show stronger intra-guild variation, with a wider range in the predictive power of climate variables relative to AM fungal genera. Genus-level data is presented here for visual clarity; taxon-level variables for all AM and EcM fungi can be found in Table S2.

The climatic variable with largest difference in observed ranges between AM and EcM fungal taxa was precipitation seasonality (47.26% greater for AM than EcM fungi, r = 0.74, p < 0.001). Soil pH and % sand content were higher for AM than EcM fungi (8.6% for sand content, r = 0.21, 19.3% for soil pH, r = 0.39, p < 0.001), though we did not find differences in range values between mycorrhizal fungal guilds for soil organic carbon (SOC) content, depth to bedrock, or soil thickness (p > 0.05). Most plant cover variables had greater ranges for AM fungi (aboveground biomass, NPP, EVI coefficient of variation, and evergreen broadleaf, shrub, herbaceous vegetation, and barren cover, r = 0.22 – 0.61, p < 0.001) while EcM fungal taxa had a greater range in evergreen needleleaf tree cover (r = 0.60, p < 0.001). Finally, AM fungi occupied a more diverse set of disturbance conditions, with over 10x larger range of burn levels (r = 0.75, p < 0.001) and 2.5x larger range of human population density levels (r = 0.42, p < 0.001) for AM fungal taxa relative to EcM fungi.

### H2 Environmental correlates of mycorrhizal fungal distributions

In support of our hypothesis that niche determinants of mycorrhizal fungal taxa varied between guilds, we found broad differences in the importance of environmental covariates for AM and EcM fungal niches. While the most important variables (based on RF % decrease in node impurity) were generally shared between AM and EcM fungi (Figure 2), such as mean annual temperature, SOC, net primary productivity (NPP), and potential evapotranspiration (PET), their relative strength differed. In support of abiotic drivers as primary determinants of EcM fungal niches, climate variables were significantly more important for explaining the occurrence of EcM fungal taxa (median = 36.0%, IQR = 5.7%) than for AM fungi (median = 30.0%, IQR = 4.6%; p < 0.001). Biotic niche drivers had lower relative feature importance than climate for EcM fungi (median 29.0%, IQR 3.6%), but similar for AM fungi (median 28.3%, IQR 2.6%). The influence of plant cover was greater for NPP in AM fungi and for evergreen need leaf tree cover in EcM fungi. Whereas the aggregate importance of soil properties was similar for both mycorrhizal guilds, though larger for AM than EcM fungi (AM median 18.9%, EcM median 16.5%, IQR 3%; p < 0.001), individual features differed, with SOC a stronger predictor for EcM fungi while sand content and soil pH more important for AM fungi (Figure 2). Arbuscular mycorrhizal fungal occurrence was also more strongly influenced by landscape physical properties (median: 17.2 ± 2.4% AM vs 14.8 ± 3.5% EcM) and disturbance (5.6 ± 2.2% AM vs 3.4 ± 1.6% EcM).

Intraguild variability in the importance of niche drivers among genera differed between AM and EcM fungi. For EcM genera, the relative importance of climate nearly doubled between the lowest (*Pisolithus*: 26%) and highest (*Leccinum*: 46%) values. In contrast, climate importance varied far less among AM genera (minimum *Pacispora*: 28%; maximum *Archaeospora*: 33%). To investigate the sources of intra-guild variability, we assessed the degree of phylogenetic signal of mycorrhizal fungal genera with the relative importance of environmental variable groups. Phylogenetic signal was evident in variable importance across EcM genera for multiple variable groups with Pagel’s λ = 0.70 (p < 0.001) for the relative importance of plant cover and λ = 0.81 (p < 0.001) for soil properties (Figure S2). In contrast, the AM genera we investigated had no significant phylogenetic structure in the relative importance of niche drivers, though a marginal branch-length scaling with disturbance importance (Blomberg’s K = 0.66, p = 0.06, Figure S2). Although for AM fungi at the VTX level, we found weak clustering with shared ancestry in relation to soil property importance (λ = 0.21, p = 0.03), there was no significant phylogenetic signal for any other variables (Figure S3).

### H3 Sensitivity to projected changes in climate

To assess potential sensitivity of mycorrhizal fungi to projected future climates, we used niche modeling to estimate current range size of their geographic distributions. Differences in niche determinants and breadth led to vastly different spatial distributions for AM and EcM fungal taxa. Based on our niche models, we estimate that the median geographic range of potentially suitable habitat for an AM fungal taxon (36 million km^2^; IQR = 24 million km^2^) is over three times larger than that of an EcM fungal taxon (11 million km^2^; IQR = 14 million km^2;^ p < 0.001; Figure 4a), or an area roughly the size of North America (∼25 million km^2^). Distributions of AM fungal taxa ranged in size between 96.1 million km^2^ (*Glomus VTX00167*) and 21.2 million km^2^ (*Acaulospora VTX00027*) while EcM fungal distributions ranged between 50.9 million km^2^ (*Tomentella subclavigera*) and 0.1 million km^2^ (*Inocybe pseudoreducta;* Table S2).

**Figure 4.**
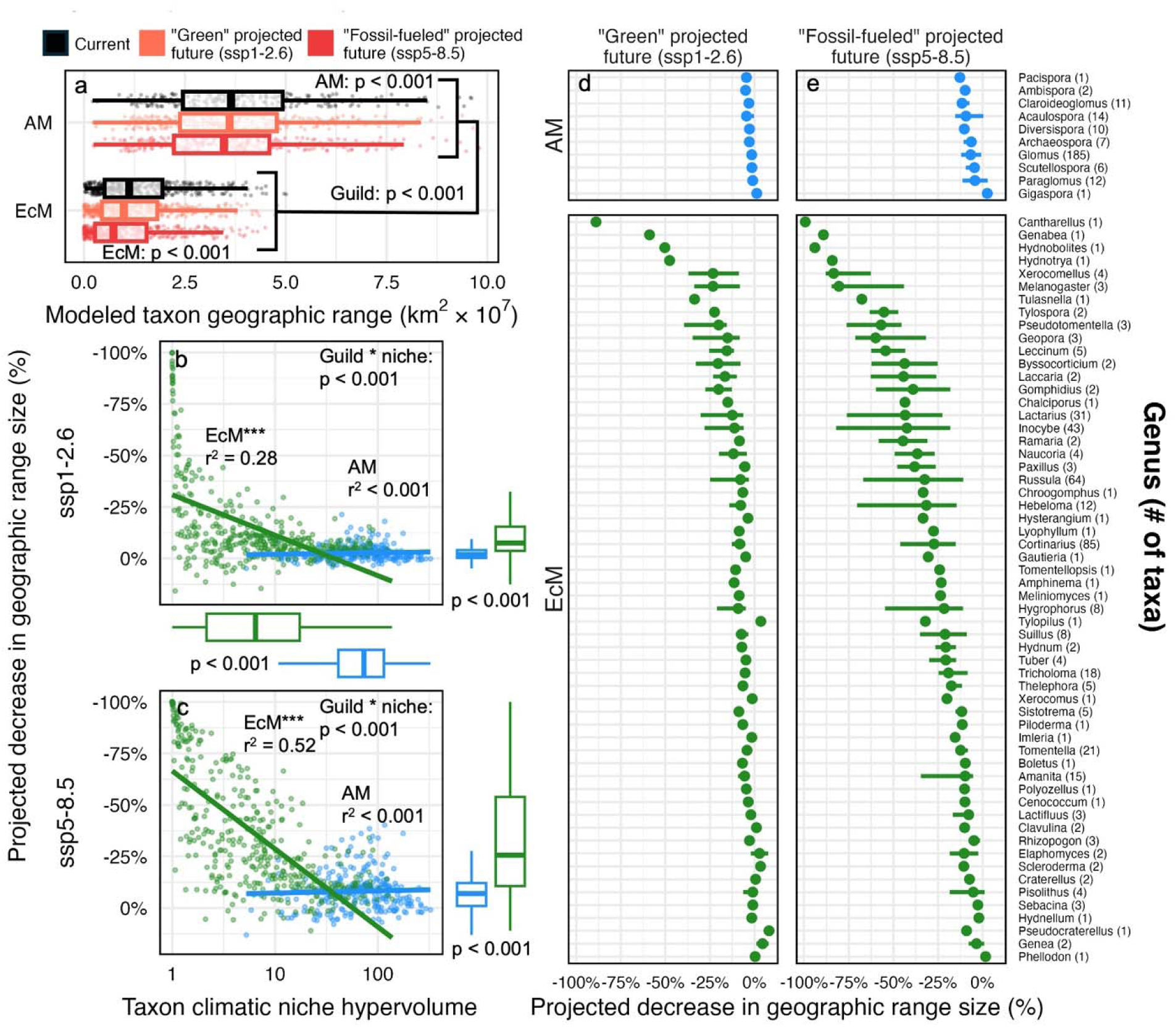
Current and projected future geographic range sizes of mycorrhizal fungal taxa (a), relationship between projected future geographic range shifts and climatic niche hypervolume (b and c), and median projected range shifts of mycorrhizal fungal genera (d and e). Range projections were generated using random forest modeling with global environmental variable estimates as predictors (Table S1). Future projections were generated similarly, but with substituting current climate data for projected future climate data from the CMIP6 models (2071 – 2100, ssp1-2.6 - sustainable “green” scenario, ssp5-8.5 - high emission “fossil fuel dependent” scenario) as predictors. Taxon range was calculated as the total area of the modeled taxon occurrence map (described further in methods and Figure S1). Guild in (a) indicates a significant difference between AM and EcM ranges, p-values within guilds indicate differences in projected range size under various climate scenarios. Similarly, in (c and d) “Guild * niche” represents a significant interaction where range contractions increase with decreasing niche hypervolumes for EcM fungi while this relationship does not persist for AM fungi. Boxplots for differences in projected range shifts and climatic niche hypervolumes between mycorrhizal fungal guilds are shown in the margins of (b and c), with p-values based on Wilcoxon rank-sum test. In (d and e) points represent the median projected range shift for each genus with error bars showing the inter-quartile range. The value next to each genus name shows the number of taxa in each genus that were analyzed. Genus-level data is presented here for visual clarity, taxon-level variables for all AM and EcM fungi can be found in Table S2.

Our models predicted widespread decreases in geographic range sizes for mycorrhizal fungal taxa based on their niche determinants under both “best” (ssp1-2.6) and “worst” (ssp5-8.5) case climate projections, with greater changes for EcM relative to AM fungi. Distributions of EcM fungi were more sensitive than AM fungi to projected future climate conditions (p < 0.001, Figure 4). Under severe simulated climate change (ssp5-8.5) median predicted decrease in range size of AM fungal taxa was 7.0 ± 5.5% (2.0 ± 1.8 million km^2^) of their current range while the median decrease for EcM fungal taxa was 25.6 ± 22.5% of their current range (2.3 ± 1.4 million km^2^). Consequently, median predicted range contractions were over double for EcM fungal taxa than AM fungal taxa in terms of proportion of their current range and 13.4% larger in terms of total area (p < 0.001, Figure 4). We found smaller changes to projected fungal ranges under less sever future climate change scenarios (ssp1-2.6) with only 1.8% area decrease for AM fungi and 7.5% decrease for EcM fungi (27% and 29% of ssp5-8.5 range shift, respectively). Interestingly, some of the taxa with the greatest and smallest predicted changes in range size belong to the same genus. For EcM fungi, these included *Inocybe* (*I. erubescens*: 99% decrease vs *I. oblectabilis*: 1.3% increase), *Cortinarius* (*C. umbrinobellus*: 88% vs *C. livido-ochraceus*: 0% decrease), and *Russula* (*R. pelargonia*: 99% vs *R. violeipes*: 3.6% decrease).

AM fungi also had large variation in predicted changes in range size across both highly diverse genera (*Glomus VTX00123*: 40.2% decrease vs *Glomus VTX00363*: 7% increase) and less diverse genera (*Acaulospora VTX00014:* 33% decrease vs *Acaulospora VTX00027*: 13% increase). Taxa with the greatest and smallest predicted range shifts are presented in Figure S4. The most vulnerable AM fungal taxa belonged to *Glomus* and *Acaulospora* genera while the most vulnerable EcM taxa belonged to *Tomentella, Inocybe, Tricholoma, Russula, Cantharellus,* and *Lactarius* genera. Despite projected range shift variation among and within AM and EcM fungal genera, we found significant phylogenetic clustering for EcM genera in their current and future range sizes (K > 0.87, p < 0.05; λ > 0.99, p < 0.001), but not for relative range shifts or any metrics for AM fungal genera. We also assessed phylogenetic signal for range shifts at the VTX level for AM fungi to account for greater relative lumping of taxa and similarly found no significant patterns (p > 0.05, Table S3).

We found a distinct response in the relationship between predicted change in range size and climate niche breadth across AM and EcM fungi under both ssp1-2.6 and ssp5-8.5 climate scenarios (Figure 4, p < 0.001). Our hypothesis that smaller niche breadths would lead to greater range shifts was supported for EcM fungi, where we saw a strong linear relationship (ssp1-2.5: r^2^ = 0.28; ssp5-8.5: r^2^ = 0.28; p < 0.001) of taxa with larger climatic niche hypervolumes experiencing lower predicted range contraction. However, this relationship disappeared for AM fungi, where we found no direct relationship between climatic niche and predicted change in range size under future climates (r^2^ = 0.001, p > 0.05). While there was no phylogenetic signal for modeled range shifts, climatic hypervolume clustered among closely related EcM fungal genera (K = 0.68, p = 0.04; λ = 0.92, p = 0.02). Phylogenetic generalized least squares (PGLS) analysis confirmed that, at the genus level, the relationship between niche hypervolume and projected range shift remained consistent when accounting for phylogenetic lineage (p < 0.001).

## Uncertainty

Several areas of uncertainty (63) arise from this approach including sampling bias, taxonomic resolution, and our modeling approach.

### Sampling bias

Our data reflect a global reality that relatively few soils and plants on earth have been characterized for mycorrhizal fungi (64). Particularly underrepresented, AM fungi are estimated to remain unsampled in >70% of ecoregions globally (65). To assess uncertainty in our models we overlaid predicted taxon occurrence probabilities with independent occurrence observations (Figure S1). The average modeled probability of a location with known taxon occurrence was >90%, supporting the validity of these distributions even in under sampled regions. Model accuracy increased in temperate/ boreal regions and those nearer to coasts while we found increased uncertainty in tropical and continental ecosystems where sampling effort was lower. We further accounted for sampling bias by restricting analyses to fungal taxa that had at least 50 occurrence observations across the globe. However, this likely also biased our results against mycorrhizal fungi with narrow niches or those occurring in less-sampled biomes (e.g., tropical forests). Undoubtedly, we have not sampled the entire niche of most mycorrhizal fungal taxa and future efforts directed at areas of high uncertainty in our distribution models could further refine knowledge of mycorrhizal fungal distributions and niche determinants (66).

### Taxonomic resolution

Differences in evolutionary history have led to the use of distinct molecular primers for identification of EcM (ITS) and AM fungi (SSU), resulting in varying levels of taxonomic resolution for EcM (species) and AM (VTX) fungi. To best utilize the most available data for mycorrhizal fungal occurrences, we assessed mycorrhizal guilds at different taxonomic levels: 18S-derived “virtual taxon” level for AM fungi (67) and ITS-derived species level for EcM fungi (68). These differences in taxonomic resolution could lead to more specific definitions for EcM than AM fungi, potentially resulting in smaller niches. Virtual taxa for AM fungi based on SSU sequences bias against certain families and can lump together groups at finer levels of taxonomic resolution (69, 70). Other regions, like LSU or functional gene specific barcodes (71, 72) could provide more accurate species identification; however, data availability precludes the use of this region is a study such as ours.

### Modeling approach

In predicting future changes in range sizes, we focused the potential effects of simulated climate change based on the high relative importance of climate for niche models and the greater availability of future climate projections relative to our other niche axis variables (73). However, this excludes other environmental changes that will occur like soil properties (74), plant cover (75), and ecosystem sensitivity to disturbance (76), as well as PET, a climate variable not simulated in our projections. While these inconsistencies may bias range estimates, as with all variables not simulated under future conditions, this bias is consistent across all taxa and climate projections. These changes in unsimulated variables will contribute to future realized niches of mycorrhizal fungi. For example, understanding future shifts in host plant species ranges could further refine distributional maps and niche estimates for mycorrhizal fungi. While many AM fungi are host generalists (77), accumulating evidence suggest some specialize in plant hosts species (78) with strong variation in plant growth benefits (79, 80). Considering how mycorrhizal symbiosis can constrain biogeographic patterns across plants and fungi (81, 82), and inclusion of other traits (like dispersal and evolutionary adaptation) will further enhance range shift projections under global change (83–85).

## Discussion

Our results demonstrate fundamental differences in the environmental factors shaping mycorrhizal fungal niche characteristics and their potential responses to global change. AM fungi exhibit broader niches, characterized by larger geographic distributions and a lower relative importance of wide-scale variables like climate in determining their ranges. Instead, AM fungi occurrence is selected for at the landscape-scale, with greater relative influence of local factors like soil characteristics, physical properties, and disturbance. In contrast, EcM fungal taxa occupy more narrow niches with a greater influence of climate factors on their distributions. These differences suggest trade-offs between generalization and specialization in the mycorrhizal fungal niche: AM fungi benefit from broader flexibility and wider range of potential distributions, while greater climate sensitivity in EcM fungi may lead to higher vulnerability under future climate regimes.

### Realized niche breadth and drivers

In line with our hypothesis, AM fungi consistently occupied more diverse environmental conditions than EcM fungi, with an average 63% greater range across all independent niche axes. These patterns are only partially reflected in the plant counterparts of these mycorrhizal fungi. Aboveground, climate is frequently the primary driving force in determining AM vs EcM plant occupancy due to its influence on nutrient cycling and decomposition (86, 87). However, when controlling for climate, EcM tree distributions are more reliant on local stochastic factors than specific soil conditions (88). Niche modeling for plants across different mycorrhizal associations shows that, while preferences for environmental conditions are shared between above- and below-ground symbionts (*i.e.,* warmer and wetter for AM plants and fungi), there were no differences in their niche breadths between AM and EcM plants (89). This disconnect is likely related to differences in the magnitude of diversity of plants versus fungi. Plants that associate with AM fungi are highly diverse (∼80% of plant species; 90) while AM fungal VTX diversity is considerably lower (∼400 taxa, 67). Individual AM fungal taxa associate with a wide range of plants across diverse environmental conditions (91), allowing AM fungal niches to surpass those of their plant host species. Greater niche space and more flexible environmental preferences could allow AM fungi to colonize wider spatial ranges than EcM fungi (33) and may reflect a potentially lower endemicity of AM taxa relative to EcM fungi across the globe (92, 93).

Niche breadth also varies considerably within guilds. In the largest AM fungal genus, *Glomus*, the taxon with the maximum climatic hypervolume was over 250% greater than the taxon with minimum. This variation reflects the considerable functional diversity (94) within this group as well as the general difficulty in resolving this non-monophyletic clade (95). Similarly, while EcM fungal niches were consistently smaller than AM fungi, diverse genera (*Russula*, *Inocybe*, *Cortinarius*) all contained taxa with climate niches >300% larger than their smallest. As the magnitude of this variation remains vast at the genus level, species specific characteristics like dispersal traits (85, 96) or other life history factors (84) may be necessary to determine mycorrhizal fungal distributions. Although evidence directly linking mycorrhizal fungal ranges with dispersal traits at the global scale is limited (33), dispersal can strongly influence community assembly across landscapes (97) and continents (98). This suggests that dispersal may play a critical role in future range shifts of mycorrhizal fungi under global change, yet this mechanism is not fully integrated into current projections. The considerably higher variation within EcM fungal niches than for AM fungi demonstrates a need for careful consideration of inter- and intra-guild differences in predicting distribution patterns for these organisms now and into the future.

### Sensitivity to projected changes in climate

Understanding the niche of mycorrhizal fungi is an important step for including these taxa in projections of global change sensitivity. As EcM fungal niches frequently correspond to colder and drier climates compared to AM fungi, EcM fungal taxa may be particularly susceptible to global warming (99–101). Tolerance to potential disturbance varies greatly among taxa in both AM and EcM fungal guilds (102–104). As climate disturbances increase in frequency and severity AM fungal taxa may be more likely to tolerate these shifts than EcM fungi. However, there is also large variation in ecological niches among EcM fungal taxa, which may allow these communities to compensate with greater tolerance to local plant and land use disturbances than for AM fungi (105, 106). Empirical evidence shows that under changing environments EcM fungal communities shift their functional composition to more generalist, less host specialized taxa (107). These patterns suggest that there is likely to be a wide range of responses to ongoing global change among taxa within mycorrhizal guilds, but that EcM fungi generally show greater potential climate sensitivity than AM fungi.

Our models predicted widespread decreases in suitable habitat and potential geographic ranges under changing climates for the majority of mycorrhizal fungal taxa we assessed. However, these models are unable to account for various fungal traits that may influence their responses to changing environmental conditions, such as dispersal, biotic interactions, or thermal adaptation. Dispersal limitations could further constrain potential fungal range shifts under changing climates as fungal taxa may not have access to all potentially suitable new habitats if their propagules cannot reach them (108). Differences between mycorrhizal guilds in dispersal capacity could potentially modify their proportional responses to changing climates (85, 109). As habitats warm, taxa with lower thermal tolerance will likely go locally extinct more quickly than those with greater tolerance (110, 111). Under these lower-competition scenarios, more tolerant taxa could potentially expand their realized niches and geographic range (112, 113). Further, warmer conditions with greater precipitation variability will create potential niche space that does not currently exist. These novel niche conditions could be suitable habitat to disturbance-tolerant fungal taxa or those with a relatively high capacity to adapt to more extreme environments (114, 115). Ultimately, predicting how interactions and adaptations will influence future geographic redistribution of fungi will require greater empirical evidence for the specific responses of fungal taxa and communities under changing environmental conditions.

## Conclusions

The mycorrhizal fungal niche determinants and predicted ranges we show here add to an ongoing effort to define the global microbiome (93, 116–119). Given the prevalence of mycorrhizal fungi in biogeochemical cycling (8, 24), plant productivity (120), invasion potential (121–123), and ecosystem response to global change (26), outlining the mycorrhizal fungal niche at the global scale helps refine forecasts of how their role in ecosystems might shift over the coming decades (124). Beyond supporting broader ecological hypotheses governing the occurrence of these organisms, this study also emphasizes the need for specific consideration of mycorrhizal fungi in conservation. Within mycorrhizal guilds (and genera), fungal taxa vary widely in the environmental factors that predict their occurrence and their potential sensitivity to changing climates. Both at the guild and taxon level, the potential drivers of fungal distributions and their vulnerability could be useful for developing more effective management interventions to protect terrestrial ecosystems.

## Methods

### Fungal occurrence data collection and curation

Georeferenced mycorrhizal fungal sequences from roots and soils (any depth) were collected from the NCBI GenBank (60) sequenced from 1998 through December 2024. To be included in this study, each record must (1) originate from a natural setting rather than greenhouse experiments or manipulative experiments, (2) include geographic coordinates and (3) be in the correct sequencing region: 18S for AM fungi and ITS2 for EcM fungi. We then supplemented this dataset with records of EcM fungi from GlobalFungi 5.0 (accessed December 2024; 62) and AM fungi from GlobalAMFungi 1.0 (accessed December 2024; 61). All sequences were grouped at ∼97% identity (V4 variable region of 18S for AM fungi and ITS2 for EcM fungi) and classified into established molecular taxa for AM fungi (virtual taxa; VTXs) or EcM fungi (Species Hypotheses; SHs) using the MaarjAM (67) and UNITE (125) databases, respectively. Taxa were confirmed to be mycorrhizal using established lists of AM and EcM fungal taxa from the MaarjAM (67) and FungalTraits (126) databases.

### Dataset description

Our initial data search resulted in 150,831 observations from 1,004 fungal taxa (625 EcM and 379 AM). We removed 128 taxa (all EM; 23,367 observations) based on conflicting reports of mycorrhizal associations in the literature, such as different guilds or lifestyles across fungal functional identification databases (126, 127). An additional 255 taxa (114 EcM and 141 AM; 6,385 observations) were removed because these taxa had fewer than 50 recorded global observations in our database. For our final analysis, we use 115,924 georeferenced samples of mycorrhizal fungal rRNA taxon occurrences (Figure 1) to map the geographic distributions and ecological niches of 249 AM (VTX) and 402 EcM fungal (SH) taxa. To ensure adequate sampling depth, we only retained those mycorrhizal fungal taxa that had more than 50 unique geographic occurrences.

### Environmental Covariate Data

Environmental covariates used for global niche modeling were chosen from a candidate set of 106 global environmental covariate layers, which describe variation in climate (128, 129), soil physiochemical characteristics (130, 131), topography (132), vegetation (133, 134), and anthropogenic influence (133). Because many of these covariates are strongly correlated, we selected a subset of less correlated, ecologically meaningful variables. To do so, we performed a principal component analysis on 100 globally distributed random terrestrial locations of all environmental covariate layers, which showed 30 principal components could explain ∼90% of the global variation in all 106 environmental covariates. We then used hierarchical clustering, which grouped environmental covariates into 31 groups of covariates most closely associated with each principal component. We selected variables from these clusters that corresponded to five main categories: climate, plant cover, physical characteristics, soil characteristics and disturbance. This resulted in 31 global environmental covariates that were used for global species distribution modeling (Table S1). Next, we summarized each of the 31 environmental layers described above for each mycorrhizal fungal taxon using a mean, median, minimum, maximum and standard deviation.

To model contemporary realized mycorrhizal fungal niches under predicted future climate conditions for 2071-2100, we used CMIP6 projections from CHELSA (129). Due to tight coupling and collinearity among soil variables, we only included soil C and pH in all models and excluded soil N or P. In addition to avoiding collinearity among soil C and N/P, SOC metrics often provide a more reliable index of soil fertility because total stocks of soil nutrients do not accurately reflect available nutrient pools, which are difficult to model given their rapid turnover (135).

### Niche Modeling and Mapping

We used the random forest machine-learning algorithm to produce niche models that correlate fungal occurrence data to environmental covariates. For each taxon, we aggregated all observations into unique pixels (at 300 arc-second resolution; 10 km^2^) before model training. We then selected an equal number of pseudo-absences that were not within 2 degrees of any focal fungal taxon presence observation (136). Combining these presences and pseudo-absences with the pixel values for our selection of 31 environmental variables, we trained a random forest algorithm to produce our models (number of trees = 100, variables per split = 5.6, minimum leaf population = 1), and generated relative occurrence probability (or relative occurrence rate) maps for each mycorrhizal fungal taxon. To reduce potential bias of the selection of pseudo-absences, this process was repeated 10 times, each time using a different random seed for generating pseudo-absences. The final spatial predictions are the mean probabilities across these 10 iterations. To generate relative occurrence probability maps for future climate scenarios, we used the random forest model trained with current climate (and other environmental) data and replaced climate bands (MAT, MAP, max temp of warmest month, and precipitation seasonality) from 2071-2100 (ssp1-2.6 and ssp5-8.5) CMIP6 projections in the CHELSA database. PET is not included in this database and was not simulated for future climate projections. Results were exported as georeferenced raster images with bands for current and scenario-specific probabilities. To assess the degree of extrapolation between training and future simulated climate data, we calculated climatic envelopes (min and max values) for each taxon and simulated climate variables. We then calculated the total proportion of future area under each climate simulation and the number of variables that fell outside these envelopes (Figure S5).

To evaluate model performance and uncertainty, we used a random cross-validation approach with a 50% probability threshold to classify each point. For each cross-validation iteration, accuracy scores and Cohen’s kappa statistics were calculated to show levels of agreement between observed and expected classifications. Model uncertainty was quantified by aggregating results across all 10 iterations. To further quantify prediction uncertainty, we performed bootstrapped resampling of the training data (based on the number of observations), retrained models on each sample, and computed ensemble means and confidence intervals for each pixel’s predicted occurrence probability. Random Forest feature importance scores were computed as the summed decrease of impurity over all trees in the forest that results from the variable used at a given node. Final feature importance scores are taken as the mean feature importance across all 10 model iterations. All computations were run on Google Earth Engine (Gorelick et al., 2017) using the Python API.

### Estimating Niche Hypervolumes

Climatic niche hypervolumes (n-dimensional space representing the full set of environmental conditions where a species can persist) were calculated with the R package *hypervolume* (137). We used the current climate predictors (Mean Annual Temperature, Maximum Temperature of Warmest Month, Precipitation Seasonality, Potential Evapotranspiration, and Mean Annual Precipitation) and modeled occurrence data. For each taxon, a maximum of 2,500 unique observations were selected and the hypervolume was calculated using Gaussian kernel density estimation.

### Statistical analyses and summaries

### Niche determinants and breadth (H1)

All statistics were conducted in R v. 4.1.0 (138). To test the hypothesis that mycorrhizal fungal guilds differ in the basic environmental conditions they occupy, mean mycorrhizal fungal climatic niches were calculated based on the mean annual temperature (MAT) and mean annual precipitation (MAP) using the CHELSA climate layers (139). Mean climatic niches for each mycorrhizal fungal taxon were overlaid on Whittaker biome maps (140) using the plotbiomes R package (141). To test the hypothesis that AM and EcM fungi differ in the breadth of their niches, we compared the total range of environmental conditions (max – min) where taxa occur for all environmental niche axes (Table S1). Differences in the median value and range of independent niche axes for AM and EcM fungi were compared using the Wilcoxon sign-rank test, due to the non-normal distribution of these data, with mycorrhizal fungal guild as the main effect. We also corrected for multiple comparisons using a Benjamini–Hochberg p-value adjustment.

### Abiotic vs biotic niche drivers (H2)

To test the hypothesis that the drivers of fungal niches differ between AM and EcM fungi, we used the relative variable importance of independent niche axes from the Random Forest model. Variables were classified into 5 different categories (climate, soil properties, plant cover, physical properties, and disturbance). Variable importance (relative % decrease in MSE) of each fungal taxon and environmental covariates was derived from global niche models at a taxon-specific accuracy threshold chosen using a decile approach where 90% of observations were captures at that threshold (Figure S1). This threshold was chosen as true-absence data was unavailable for generating and validating these organismal distribution models. The relative importance of each variable within a category was summed for each taxon. Differences between mycorrhizal fungal guilds in mean category importance values were tested using the same approach described for H1 (Wilcoxon sign-rank and BH adjustment). We also assessed phylogenetic signals for niche drivers among AM and EcM fungal genera as well as AM fungal VTX by assessing Blomberg’s K and Pagel’s λ using the phylosig() function in the phytools 2.0 package (142). Phylogenetic trees used for these analyses were generated according to Kivlin 2020 and are available in the supplementary information.

### Sensitivity to global change (H3)

To test the hypothesis that mycorrhizal fungal taxa with narrower niches will be more sensitive to projected changes in climate, we compared the relationship of projected range shifts with climatic niche hypervolumes. Current and projected future range sizes of each taxon were estimated using the modeled occurrence probability maps generated from the previously described niche modeling/mapping. Differences between mycorrhizal fungal guilds in their range shifts were assessed using 1) a mixed effect linear model with range size as the response variable and mycorrhizal fungal guild, climate scenario (current, ssp1-2.6, ssp5-8.5), and their interaction as fixed effects and mycorrhizal taxon as a random effect; and 2) assessing the % change in range size within each mycorrhizal fungal taxon and comparing range shifts between AM and EcM fungal guilds using a Wilcoxon sign-rank test. We also used this test to assess differences between mycorrhizal fungal guilds in their climatic niche hypervolume. To assess whether that relationship between climatic niche hypervolume and projected range shift differed between mycorrhizal fungal guilds, we used a linear model with projected range shift as the response variable and mycorrhizal fungal guilds, niche hypervolume, and their interactions and fixed effects. We also assessed phylogenetic signals for climatic niche hypervolume, current range size, future projected range size, and projected range shift among AM and EcM fungal genera as well as AM fungal VTX by assessing Blomberg’s K and Pagel’s λ as previously described. Based on significant clustering with shared ancestry for EcM fungi of climatic niche hypervolumes, we confirmed the correlations with projected range shift after accounting for phylogeny via PGLS regression.

## Acknowledgements

JDE was supported by NSF award 2305863. SNK was funded by NSF awards 2106065, 2338421, and 2412561 as well as start-up funding from the University of Tennessee. The authors thank the Kivlin, Bailey, and Schweitzer laboratory groups, particularly Ella Segal, for thoughtful conversations about this manuscript. PK was supported from the Czech Science Foundation project: The effect of global changes on fungal biogeography and ecosystem functioning (21-20802M) and the Ministry of Education, Youth and Sports of the Czech Republic grant Talking microbes - understanding microbial interactions within One Health framework (CZ.02.01.01/00/22_008/0004597). JDS, CQ, MVN, and ETK were funded with grants from the Jeremy and Hannelore Grantham Environmental Trust, the Paul Allen Family Foundation, Bezos Earth Fund, Schmidt Family Foundation, Hefner Foundation. JDS and ETK were funded by the NWO Gravity Grant MICROP [024.004.014]. ETK was also supported by a NWO-VICI [202.012], HFSP [RGP 0029], NWO-Spinoza [SP.2023.2] (ETK), and an Ammodo grant (ETK). CSD was supported by a Swiss National Science Foundation Postdoctoral Fellowship (TMPFP3_209925).

## References

1. W. Thuiller, S. Lavorel, M. B. Araújo, Niche properties and geographical extent as predictors of species sensitivity to climate change. Global Ecol Biogeogr 14, 347–357 (2005).

2. M. Gerz, C. Guillermo Bueno, W. A. Ozinga, M. Zobel, M. Moora, Niche differentiation and expansion of plant species are associated with mycorrhizal symbiosis. Journal of Ecology 106, 254–264 (2018).

3. A. Mestre et al., The interplay of nested biotic interactions and the abiotic environment regulates populations of a hypersymbiont. Journal of Animal Ecology 88, 1998–2010 (2019).

4. K. G. Peay, The mutualistic niche: mycorrhizal symbiosis and community dynamics. Annual Review of Ecology, Evolution, and Systematics 47, 143–164 (2016).

5. R. J. Smith, S. Jovan, B. McCune, Climatic niche limits and community-level vulnerability of obligate symbioses. Journal of Biogeography 47, 382–395 (2020).

6. M. G. van Der Heijden, F. M. Martin, M. A. Selosse, I. R. Sanders, Mycorrhizal ecology and evolution: the past, the present, and the future. New Phytol 205, 1406–1423 (2015).

7. C. Averill et al., Defending Earth’s terrestrial microbiome. Nature Microbiology 7, 1717–1725 (2022).

8. H.-J. Hawkins et al., Mycorrhizal mycelium as a global carbon pool. Current Biology 33, R560–R573 (2023).

9. S. W. Behie, M. J. Bidochka, Nutrient transfer in plant–fungal symbioses. Trends in plant science 19, 734–740 (2014).

10. M. E. Van Nuland et al., Global hotspots of mycorrhizal fungal richness are poorly protected. Nature, 1–9 (2025).

11. L. Tedersoo, M. Bahram, M. Zobel, How mycorrhizal associations drive plant population and community biology. Science 367, eaba1223 (2020).

12. B. M. Ohsowski, P. D. Zaitsoff, M. Öpik, M. M. Hart, Where the wild things are: looking for uncultured Glomeromycota. New Phytol 204, 171–179 (2014).

13. F. Martin, A. Kohler, C. Murat, C. Veneault-Fourrey, D. S. Hibbett, Unearthing the roots of ectomycorrhizal symbioses. Nature Reviews Microbiology 14, 760–773 (2016).

14. L. Tedersoo, M. E. Smith, Lineages of ectomycorrhizal fungi revisited: foraging strategies and novel lineages revealed by sequences from belowground. Fungal biology reviews 27, 83–99 (2013).

15. O. Comandini, A. C. Rinaldi, T. W. Kuyper, Measuring and estimating ectomycorrhizal fungal diversity: a continuous challenge. Mycorrhiza: occurrence in natural and restored environments, 165–200 (2012).

16. D. Janowski, T. Leski, Methods for identifying and measuring the diversity of ectomycorrhizal fungi. Forestry: An International Journal of Forest Research 96, 639–652 (2023).

17. L. G. Van Galen et al., The biogeography and conservation of Earth’s ‘dark’ectomycorrhizal fungi. Current Biology 35, R563–R574 (2025).

18. M. C. Brundrett, L. Tedersoo, Evolutionary history of mycorrhizal symbioses and global host plant diversity. New Phytol 220, 1108–1115 (2018).

19. S. Miyauchi et al., Large-scale genome sequencing of mycorrhizal fungi provides insights into the early evolution of symbiotic traits. Nat Commun 11, 5125 (2020).

20. L. Tedersoo, T. W. May, M. E. Smith, Ectomycorrhizal lifestyle in fungi: global diversity, distribution, and evolution of phylogenetic lineages. Mycorrhiza 20, 217–263 (2010).

21. C. Averill, J. M. Bhatnagar, M. C. Dietze, W. D. Pearse, S. N. Kivlin, Global imprint of mycorrhizal fungi on whole-plant nutrient economics. Proceedings of the National Academy of Sciences 116, 23163–23168 (2019).

22. M. E. Craig et al., Tree mycorrhizal type predicts within-site variability in the storage and distribution of soil organic matter. Global Change Biol 24, 3317–3330 (2018).

23. G. Lin, M. L. McCormack, C. Ma, D. Guo, Similar below-ground carbon cycling dynamics but contrasting modes of nitrogen cycling between arbuscular mycorrhizal and ectomycorrhizal forests. New Phytol 213, 1440–1451 (2017).

24. R. P. Phillips, E. Brzostek, M. G. Midgley, The mycorrhizal-associated nutrient economy: a new framework for predicting carbon-nutrient couplings in temperate forests. New Phytol 199, 41–51 (2013).

25. C. N. Barnes et al., Root causes of below-ground processes: Tree mycorrhizal associations outweigh tree richness. Journal of Ecology 114, e70282 (2026).

26. C. Averill, M. C. Dietze, J. M. Bhatnagar, Continental-scale nitrogen pollution is shifting forest mycorrhizal associations and soil carbon stocks. Global Change Biol 24, 4544–4553 (2018).

27. P. Baldrian, R. López-Mondéjar, P. Kohout, Forest microbiome and global change. Nature Reviews Microbiology 21, 487–501 (2023).

28. I. Jo, S. Fei, C. M. Oswalt, G. M. Domke, R. P. Phillips, Shifts in dominant tree mycorrhizal associations in response to anthropogenic impacts. Science advances 5, eaav6358 (2019).

29. R. Liese, C. Leuschner, I. C. Meier, The effect of drought and season on root life span in temperate arbuscular mycorrhizal and ectomycorrhizal tree species. Journal of Ecology 107, 2226–2239 (2019).

30. A. F. Pellegrini et al., Decadal changes in fire frequencies shift tree communities and functional traits. Nature Ecology & Evolution 5, 504–512 (2021).

31. C. Terrer et al., A trade-off between plant and soil carbon storage under elevated CO2. Nature 591, 599–603 (2021).

32. E. C. Segal, S. N. Kivlin, Determinants of Plant–Mycorrhizal Fungal Distributions and Function Under Global Change. Annual Review of Ecology, Evolution, and Systematics 56 (2025).

33. S. N. Kivlin, Global mycorrhizal fungal range sizes vary within and among mycorrhizal guilds but are not correlated with dispersal traits. Journal of Biogeography 47, 1994–2001 (2020).

34. J. D. Hoeksema, Investigating the disparity in host specificity between AM and EM fungi: lessons from theory and better-studied systems. Oikos 84, 327–332 (1999).

35. F. Voller, A. Ardanuy, A. F. Taylor, D. Johnson, Maintenance of host specialisation gradients in ectomycorrhizal symbionts. New Phytol 242, 1426–1435 (2024).

36. J. D. Edwards, J. W. Dalling, J. M. Fraterrigo, W. C. Eddy, W. H. Yang, Functional traits of ectomycorrhizal trees influence surrounding soil organic matter properties. Functional Ecology (2025).

37. J. Davison et al., Temperature and pH define the realised niche space of arbuscular mycorrhizal fungi. New Phytol 231, 763–776 (2021).

38. Y. Miyamoto, Y. Terashima, K. Nara, Temperature niche position and breadth of ectomycorrhizal fungi: reduced diversity under warming predicted by a nested community structure. Global Change Biol 24, 5724–5737 (2018).

39. C. Qin, P. T. Pellitier, M. E. Van Nuland, K. G. Peay, K. Zhu, Niche modelling predicts that soil fungi occupy a precarious climate in boreal forests. Global Ecol Biogeogr 32, 1127–1139 (2023).

40. T. Větrovský et al., A meta-analysis of global fungal distribution reveals climate-driven patterns. Nat Commun 10, 5142 (2019).

41. I. Hiiesalu et al., Latitudinal and Elevational Trends in Arbuscular Mycorrhizal Community Niche Structure. Ecol Lett 28, e70241 (2025).

42. M. Buée, P.-E. Courty, D. Mignot, J. Garbaye, Soil niche effect on species diversity and catabolic activities in an ectomycorrhizal fungal community. Soil Biology and Biochemistry 39, 1947–1955 (2007).

43. S. N. Kivlin, C. V. Hawkes, K. K. Treseder, Global diversity and distribution of arbuscular mycorrhizal fungi. Soil Biology and Biochemistry 43, 2294–2303 (2011).

44. L. M. Suz et al., Environmental drivers of ectomycorrhizal communities in Europe’s temperate oak forests. Molecular Ecology 23, 5628–5644 (2014).

45. S. van der Linde et al., Environment and host as large-scale controls of ectomycorrhizal fungi. Nature 558, 243-+ (2018).

46. T. M. Mansfield, F. E. Albornoz, M. H. Ryan, G. D. Bending, R. J. Standish, Niche differentiation of Mucoromycotinian and Glomeromycotinian arbuscular mycorrhizal fungi along a 2-million-year soil chronosequence. Mycorrhiza 33, 139–152 (2023).

47. M. E. Van Nuland, K. G. Peay, Symbiotic niche mapping reveals functional specialization by two ectomycorrhizal fungi that expands the host plant niche. Fungal Ecology 46, 100960 (2020).

48. J. M. Chase, M. A. Leibold, Ecological niches: linking classical and contemporary approaches (University of Chicago Press, 2009).

49. D. Nogués-Bravo, Predicting the past distribution of species climatic niches. Global Ecol Biogeogr 18, 521–531 (2009).

50. A. E. Bennett, A. T. Classen, Climate change influences mycorrhizal fungal–plant interactions, but conclusions are limited by geographical study bias. Ecology 101, e02978 (2020).

51. J. D. Edwards et al., Warming disrupts Plant–Fungal endophyte symbiosis more severely in leaves than roots. Global Change Biol 31, e70207 (2025).

52. M. Qi et al., Predicted Effects of Climate Change on Future Distributions of Ectomycorrhizal Fungi. Ecology and Evolution 15, e72743 (2025).

53. B. Blonder, C. Lamanna, C. Violle, B. J. Enquist, The n-dimensional hypervolume. Global Ecol Biogeogr 23, 595–609 (2014).

54. M. Chevalier, O. Broennimann, A. Guisan, Climate change may reveal currently unavailable parts of species’ ecological niches. Nature Ecology & Evolution 8, 1298–1310 (2024).

55. S. J. Davies, M. P. Hill, M. A. McGeoch, S. Clusella-Trullas, Niche shift and resource supplementation facilitate an amphibian range expansion. Diversity and Distributions 25, 154–165 (2019).

56. N. A. Freymueller, E. D. Lorenzen, S. C. Brown, C. Rahbek, D. A. Fordham, 21st Century Sea Ice Loss Will Upend 11,700 Years of Stable Habitat for Bowhead Whales. Ecology and Evolution 15, e71377 (2025).

57. P. C. Le Roux, M. A. McGeoch, Rapid range expansion and community reorganization in response to warming. Global Change Biol 14, 2950–2962 (2008).

58. T. M. Chaloner, S. J. Gurr, D. P. Bebber, Geometry and evolution of the ecological niche in plant-associated microbes. Nat Commun 11, 2955 (2020).

59. M. E. Van Nuland, C. Qin, P. T. Pellitier, K. Zhu, K. G. Peay, Climate mismatches with ectomycorrhizal fungi contribute to migration lag in North American tree range shifts. Proceedings of the National Academy of Sciences 121, e2308811121 (2024).

60. E. W. Sayers et al., Database resources of the National Center for Biotechnology Information in 2025. Nucleic Acids Res 53, D20–D29 (2025).

61. T. Větrovský et al., GlobalAMFungi: a global database of arbuscular mycorrhizal fungal occurrences from high-throughput sequencing metabarcoding studies. New Phytol 240, 2151–2163 (2023).

62. T. Vetrovsky et al., GlobalFungi, a global database of fungal occurrences from high-throughput-sequencing metabarcoding studies. Scientific Data 7 (2020).

63. J. Jansen et al., Stop ignoring map uncertainty in biodiversity science and conservation policy. Nature Ecology & Evolution 6, 828–829 (2022).

64. C. A. Guerra et al., Blind spots in global soil biodiversity and ecosystem function research. Nat Commun 11, 3870 (2020).

65. J. D. Stewart et al., Advancing knowledge on the biogeography of arbuscular mycorrhizal fungi to support Sustainable Development Goal 15: Life on Land. FEMS Microbiology Letters 372, fnaf055 (2025).

66. D. J. Baker, I. M. D. Maclean, M. Goodall, K. J. Gaston, Correlations between spatial sampling biases and environmental niches affect species distribution models. Global Ecol Biogeogr 31, 1038–1050 (2022).

67. M. Öpik et al., The online database MaarjAM reveals global and ecosystemic distribution patterns in arbuscular mycorrhizal fungi (Glomeromycota). New Phytol 188, 223–241 (2010).

68. K. Abarenkov et al., The UNITE database for molecular identification and taxonomic communication of fungi and other eukaryotes: sequences, taxa and classifications reconsidered. Nucleic Acids Res 52, D791–D797 (2024).

69. T. D. Bruns, N. Corradi, D. Redecker, J. W. Taylor, M. Öpik, Glomeromycotina: what is a species and why should we care? New Phytol 220, 963–967 (2018).

70. T. D. Bruns, J. W. Taylor, Comment on “Global assessment of arbuscular mycorrhizal fungus diversity reveals very low endemism”. Science 351, 826–826 (2016).

71. C. S. Delavaux, R. J. Ramos, S. L. Sturmer, J. D. Bever, Environmental identification of arbuscular mycorrhizal fungi using the LSU rDNA gene region: an expanded database and improved pipeline. Mycorrhiza 32, 145–153 (2022).

72. F. Stefani et al., The pitfalls of r DNA-based AMF identification: a comparative analysis of r DNA and protein-coding genes. New Phytol 248, 1501–1515 (2025).

73. B. C. O’Neill, et al., The Scenario Model Intercomparison Project (ScenarioMIP) for CMIP6. (2016).

74. D. Beillouin et al., A global overview of studies about land management, land-use change, and climate change effects on soil organic carbon. Global Change Biol 28, 1690–1702 (2022).

75. A. E. Kelly, M. L. Goulden, Rapid shifts in plant distribution with recent climate change. Proceedings of the national academy of sciences 105, 11823–11826 (2008).

76. G. Kröel-Dulay et al., Increased sensitivity to climate change in disturbed ecosystems. Nat Commun 6, 6682 (2015).

77. T. W. d’Entremont, S. N. Kivlin, Specificity in plant-mycorrhizal fungal relationships: prevalence, parameterization, and prospects. Frontiers in Plant Science 14, 1260286 (2023).

78. V. Kokkoris et al., Codependency between plant and arbuscular mycorrhizal fungal communities: what is the evidence? New Phytol 228, 828–838 (2020).

79. J. N. Klironomos, Variation in plant response to native and exotic arbuscular mycorrhizal fungi. Ecology 84, 2292–2301 (2003).

80. J. D. Hoeksema et al., A meta-analysis of context-dependency in plant response to inoculation with mycorrhizal fungi. Ecol Lett 13, 394–407 (2010).

81. C. S. Delavaux, T. W. Crowther, J. D. Bever, P. Weigelt, E. M. Gora, Mutualisms weaken the latitudinal diversity gradient among oceanic islands. Nature 627, 335–339 (2024).

82. C. S. Delavaux et al., Mycorrhizal types influence island biogeography of plants. Communications Biology 4, 1128 (2021).

83. A. Bush et al., Incorporating evolutionary adaptation in species distribution modelling reduces projected vulnerability to climate change. Ecol Lett 19, 1468–1478 (2016).

84. V. B. Chaudhary et al., What are mycorrhizal traits? Trends in Ecology & Evolution 37, 573–581 (2022).

85. S. Pehim Limbu et al., Climate-linked biogeography of mycorrhizal fungal spore traits. Proceedings of the National Academy of Sciences 122, e2505059122 (2025).

86. N. A. Soudzilovskaia et al., Global mycorrhizal plant distribution linked to terrestrial carbon stocks. Nat Commun 10, 5077 (2019).

87. B. S. Steidinger et al., Climatic controls of decomposition drive the global biogeography of forest-tree symbioses. Nature 569, 404–408 (2019).

88. J. A. Medina-Vega et al., Tropical tree ectomycorrhiza are distributed independently of soil nutrients. Nature Ecology & Evolution 8, 400–410 (2024).

89. I. Rog, D. Lerner, S. F. Bender, M. G. van Der Heijden, The increased environmental niche of dual-mycorrhizal woody species. Ecol Lett 28, e70132 (2025).

90. N. A. Soudzilovskaia et al., FungalRoot: global online database of plant mycorrhizal associations. New Phytol 227, 955–966 (2020).

91. J. Klironomos, Host-specificity and functional diversity among arbuscular mycorrhizal fungi. Microbial biosystems: New frontiers 1, 845–851 (2000).

92. J. Davison et al., Global assessment of arbuscular mycorrhizal fungus diversity reveals very low endemism. Science 349, 970–973 (2015).

93. L. Tedersoo et al., Global patterns in endemicity and vulnerability of soil fungi. Global Change Biol 28, 6696–6710 (2022).

94. T. Helgason et al., Selectivity and functional diversity in arbuscular mycorrhizas of co-occurring fungi and plants from a temperate deciduous woodland. Journal of Ecology 90, 371–384 (2002).

95. D. Schwarzott, C. Walker, A. Schüßler, Glomus, the largest genus of the arbuscular mycorrhizal fungi (Glomales), is nonmonophyletic. Molecular Phylogenetics and Evolution 21, 190–197 (2001).

96. K. G. Peay, M. G. Schubert, N. H. Nguyen, T. D. Bruns, Measuring ectomycorrhizal fungal dispersal: macroecological patterns driven by microscopic propagules. Molecular ecology 21, 4122–4136 (2012).

97. A. G. Tipton et al., Arbuscular mycorrhizal fungi taxa show variable patterns of micro-scale dispersal in prairie restorations. Frontiers in Microbiology 13, 827293 (2022).

98. P. T. Pellitier et al., Wind patterns influence the dispersal and assembly of North American soil fungal communities. Ecol Lett 28, e70130 (2025).

99. M. A. Anthony, M. Knorr, J. A. Moore, M. Simpson, S. D. Frey, Fungal community and functional responses to soil warming are greater than for soil nitrogen enrichment. Elem Sci Anth 9, 000059 (2021).

100. C. Castaño et al., Ectomycorrhizal fungi with hydrophobic mycelia and rhizomorphs dominate in young pine trees surviving experimental drought stress. Soil Biology and Biochemistry 178, 108932 (2023).

101. B. S. Steidinger et al., Ectomycorrhizal fungal diversity predicted to substantially decline due to climate changes in North American Pinaceae forests. Journal of biogeography 47, 772–782 (2020).

102. S.-K. Sepp, T. Jairus, M. Vasar, M. Zobel, M. Öpik, Effects of land use on arbuscular mycorrhizal fungal communities in Estonia. Mycorrhiza 28, 259–268 (2018).

103. C. S. Delavaux et al., Uncovering diversity within the Glomeromycota: novel clades, family distributions, and land use sensitivity. Ecology and Evolution 15, e70597 (2025).

104. B. D. Lindahl, K. E. Clemmensen, J. Stendahl, A. Dahlberg, Long-term effects of clear-cutting forestry on ectomycorrhizal fungi in boreal forest. New Phytol (2026).

105. J. Kranabetter, S. Haeussler, C. Wood, Vulnerability of boreal indicators (ground-dwelling beetles, understory plants and ectomycorrhizal fungi) to severe forest soil disturbance. Forest Ecology and Management 402, 213–222 (2017).

106. M. Pärtel et al., Historical biome distribution and recent human disturbance shape the diversity of arbuscular mycorrhizal fungi. New Phytol 216, 227–238 (2017).

107. C. W. Fernandez et al., Climate change–induced stress disrupts ectomycorrhizal interaction networks at the boreal–temperate ecotone. Proceedings of the National Academy of Sciences 120, e2221619120 (2023).

108. A. Alzate, R. E. Onstein, Understanding the relationship between dispersal and range size. Ecol Lett 25, 2303–2323 (2022).

109. C. A. Aguilar-Trigueros et al., Symbiotic status alters fungal eco-evolutionary offspring trajectories. Ecol Lett 26, 1523–1534 (2023).

110. T. Kipfer, S. Egli, J. Ghazoul, B. Moser, T. Wohlgemuth, Susceptibility of ectomycorrhizal fungi to soil heating. Fungal biology 114, 467–472 (2010).

111. J. M. Sunday, A. E. Bates, N. K. Dulvy, Thermal tolerance and the global redistribution of animals. Nature climate change 2, 686–690 (2012).

112. A. Best, K. Johst, T. Münkemüller, J. Travis, Which species will succesfully track climate change? The influence of intraspecific competition and density dependent dispersal on range shifting dynamics. Oikos 116, 1531–1539 (2007).

113. A. Ettinger, J. HilleRisLambers, Competition and facilitation may lead to asymmetric range shift dynamics with climate change. Global Change Biol 23, 3921–3933 (2017).

114. A. Bazzicalupo, Local adaptation in fungi. FEMS Microbiology Reviews 46, fuac026 (2022).

115. M. A. Naranjo-Ortiz, T. Gabaldón, Fungal evolution: major ecological adaptations and evolutionary transitions. Biological Reviews 94, 1443–1476 (2019).

116. M. Delgado-Baquerizo et al., A global atlas of the dominant bacteria found in soil. Science 359, 320-+ (2018).

117. E. Egidi et al., A few Ascomycota taxa dominate soil fungal communities worldwide. Nat Commun 10 (2019).

118. L. Tedersoo et al., Global diversity and geography of soil fungi. Science 346, 1256688 (2014).

119. M. Vasar et al., Global soil microbiomes: A new frontline of biome-ecology research. Global Ecol Biogeogr 31, 1120–1132 (2022).

120. M. A. Anthony et al., Forest tree growth is linked to mycorrhizal fungal composition and function across Europe. The ISME journal 16, 1327–1336 (2022).

121. I. A. Dickie et al., The emerging science of linked plant–fungal invasions. New Phytol 215, 1314–1332 (2017).

122. I. S. Jo, K. M. Potter, G. M. Domke, S. L. Fei, Dominant forest tree mycorrhizal type mediates understory plant invasions. Ecol Lett 21, 217–224 (2018).

123. A. Pringle et al., Mycorrhizal symbioses and plant invasions. Annual Review of Ecology, Evolution, and Systematics 40, 699–715 (2009).

124. S. Lavergne, M. E. Evans, I. J. Burfield, F. Jiguet, W. Thuiller, Are species’ responses to global change predicted by past niche evolution? Philosophical Transactions of the Royal Society B: Biological Sciences 368, 20120091 (2013).

125. R. H. Nilsson et al., The UNITE database for molecular identification of fungi: handling dark taxa and parallel taxonomic classifications. Nucleic Acids Res 47, D259–D264 (2019).

126. S. Põlme et al., FungalTraits: a user-friendly traits database of fungi and fungus-like stramenopiles. Fungal diversity 105, 1–16 (2020).

127. N. H. Nguyen et al., FUNGuild: an open annotation tool for parsing fungal community datasets by ecological guild. Fungal ecology 20, 241–248 (2016).

128. A. Trabucco, R. J. Zomer, Global aridity index and potential evapotranspiration (ET0) climate database v2. (No Title) (2018).

129. D. N. Karger et al., Climatologies at high resolution for the earth’s land surface areas. Scientific data 4, 1–20 (2017).

130. J. D. Pelletier et al., A gridded global data set of soil, intact regolith, and sedimentary deposit thicknesses for regional and global land surface modeling. Journal of Advances in Modeling Earth Systems 8, 41–65 (2016).

131. T. Hengl et al., SoilGrids250m: Global gridded soil information based on machine learning. Plos One 12, e0169748 (2017).

132. G. Amatulli et al., A suite of global, cross-scale topographic variables for environmental and biodiversity modeling. Scientific data 5, 180040 (2018).

133. M. N. Tuanmu, W. Jetz, A global 1-km consensus land-cover product for biodiversity and ecosystem modelling. Global Ecol Biogeogr 23, 1031–1045 (2014).

134. M. Santoro, GlobBiomass-global datasets of forest biomass. (No Title) (2018).

135. X. He et al., Global patterns and drivers of soil total phosphorus concentration. Earth System Science Data Discussions 2021, 1–21 (2021).

136. M. Barbet-Massin, F. Jiguet, C. H. Albert, W. Thuiller, Selecting pseudo-absences for species distribution models: How, where and how many? Methods in ecology and evolution 3, 327–338 (2012).

137. B. Blonder, D. J. Harris, hypervolume: High dimensional geometry and set operations using kernel density estimation, support vector machines, and convex hulls. R package version 2 (2018).

138. R. R Core Team, R: A language and environment for statistical computing. (2021).

139. D. N. Karger, A. M. Wilson, C. Mahony, N. E. Zimmermann, W. Jetz, Global daily 1 km land surface precipitation based on cloud cover-informed downscaling. Scientific Data 8, 307 (2021).

140. R. E. Ricklefs, The economy of nature (Macmillan, 2008).

141. V. Stefan, S. Levin, Plotbiomes: Plot Whittaker biomes with ggplot2. R package version 0.0. 0.9001 (2018).

142. L. J. Revell, phytools 2.0: an updated R ecosystem for phylogenetic comparative methods (and other things). PeerJ 12, e16505 (2024).

