## Supplementary information for "Niche constraints drive differences between mycorrhizal fungal guilds in future range shifts"

Supplementary Materials

| **Class** | **Variable** | **Citation** |
| --- | --- | --- |
| Climate | CGIAR_PET | Trabucco, A, and RJ Zomer. 2018. Global Aridity Index and Potential Evapo-Transpiration (ET0) Climate Database v2. CGIAR Consortium for Spatial Information(CGIAR-CSI). Published online, available from the CGIAR-CSI  GeoPortal at https://cgiarcsi.community |
| Climate | CHELSA_BIO_Annual_Mean_  Temperature | Karger, DN, O Conrad, J Bohner, T Kawohl, H Kreft, RW Soria-Auza, NE Zimmerman, HP Linder, and M Kessler. 2017. Climatologies at high resolution for the earth’s land surface areas. Scientific Data 4:170122. |
| Climate | CHELSA_BIO_Annual_Precipitation | Karger, DN, O Conrad, J Bohner, T Kawohl, H Kreft, RW Soria-Auza, NE Zimmerman, HP Linder, and M Kessler. 2017. Climatologies at high resolution for the earth’s land surface areas. Scientific Data 4:170122. |
| Climate | CHELSA_BIO_Max_Temperature_of_Warmest_Month | Karger, DN, O Conrad, J Bohner, T Kawohl, H Kreft, RW Soria-Auza, NE Zimmerman, HP Linder, and M Kessler. 2017. Climatologies at high resolution for the earth’s land surface areas. Scientific Data 4:170122. |
| Climate | CHELSA_BIO_Precipitation_  Seasonality | Karger, DN, O Conrad, J Bohner, T Kawohl, H Kreft, RW Soria-Auza, NE Zimmerman, HP Linder, and M Kessler. 2017. Climatologies at high resolution for the earth’s land surface areas. Scientific Data 4:170122. |
| Plant Cover | ConsensusLandCoverClass_Barren | Tuanmu, M-N, and W Jetz. 2014. A global 1-km consensus land-cover product for biodiversity and ecosystem modelling. Global Ecology and Biogeography 23:1031-1045. |
| Plant Cover | ConsensusLandCoverClass_Deciduous_Broadleaf_Trees | Tuanmu, M-N, and W Jetz. 2014. A global 1-km consensus land-cover product for biodiversity and ecosystem modelling. Global Ecology and Biogeography 23:1031-1045. |
| Plant Cover | ConsensusLandCoverClass_Evergreen_Broadleaf_Trees | Tuanmu, M-N, and W Jetz. 2014. A global 1-km consensus land-cover product for biodiversity and ecosystem modelling. Global Ecology and Biogeography 23:1031-1045. |
| Plant Cover | ConsensusLandCoverClass_Evergreen_Deciduous_Needleleaf_Trees | Tuanmu, M-N, and W Jetz. 2014. A global 1-km consensus land-cover product for biodiversity and ecosystem modelling. Global Ecology and Biogeography 23:1031-1045. |
| Plant Cover | ConsensusLandCoverClass_  Herbaceous_Vegetation | Tuanmu, M-N, and W Jetz. 2014. A global 1-km consensus land-cover product for biodiversity and ecosystem modelling. Global Ecology and Biogeography 23:1031-1045. |
| Plant Cover | ConsensusLandCoverClass_Mixed_  Other_Trees | Tuanmu, M-N, and W Jetz. 2014. A global 1-km consensus land-cover product for biodiversity and ecosystem modelling. Global Ecology and Biogeography 23:1031-1045. |
| Plant Cover | ConsensusLandCoverClass_Shrubs | Tuanmu, M-N, and W Jetz. 2014. A global 1-km consensus land-cover product for biodiversity and ecosystem modelling. Global Ecology and Biogeography 23:1031-1045. |
| Plant Cover | GlobBiomass_AboveGroundBiomass | Santoro, M. 2018. GlobBiomass - global datasets of forest biomass. PANGAEA https://doi.org./10.1594/PANGAEA.894711. |
| Plant Cover | GlobPermafrost_PermafrostExtent | Obu, J, S Westermann, A Bartsch, N Berdnikov, HH Christiansen, A Dashtseren, R Delaloye, B Elberling, B Etzelmuller, A Kholodov, A Khomutov, A Kaab, MO Leibman, AG Lewkowicz, SK Panda, V Romanovsky, RG Way, A Westergaard-Nielsen, T Wu, J Yamkhin, and D Zou. 2019. Northern hemisphere permafrost map based on TTOP modelling for 2000-2016 at 1km2 scale. Earth-Science Reviews 193:299-316. |
| Plant Cover | MODIS_NPP | MOD17A3.055: Terra Net Primary Production yearly global 1km. maintained by the NASA EOSDIS Land Processes Distributed Active Archive Center (LP DAAC) at the USGS Earth Resources Observation and Science (EROS) Center, Sioux Falls, South Dakota. |
| Physical Characteristics | EarthEnvTexture_CoOfVar_EVI | Amatulli, G, S Domisch, M-N Tuanmu, B Parmentier, A Ranipeta, J Malczyk, and W Jetz. 2018. A suite of global, cross-scale topographic variables for environmental and biodiversity mapping. Scientific Data 5:180040. |
| Physical Characteristics | EarthEnvTexture_Correlation_EVI | Amatulli, G, S Domisch, M-N Tuanmu, B Parmentier, A Ranipeta, J Malczyk, and W Jetz. 2018. A suite of global, cross-scale topographic variables for environmental and biodiversity mapping. Scientific Data 5:180040. |
| Physical Characteristics | EarthEnvTexture_Homogeneity_EVI | Amatulli, G, S Domisch, M-N Tuanmu, B Parmentier, A Ranipeta, J Malczyk, and W Jetz. 2018. A suite of global, cross-scale topographic variables for environmental and biodiversity mapping. Scientific Data 5:180040. |
| Physical Characteristics | EarthEnvTopoMed_AspectCosine | Amatulli, G, S Domisch, M-N Tuanmu, B Parmentier, A Ranipeta, J Malczyk, and W Jetz. 2018. A suite of global, cross-scale topographic variables for environmental and biodiversity mapping. Scientific Data 5:180040. |
| Physical Characteristics | EarthEnvTopoMed_AspectSine | Amatulli, G, S Domisch, M-N Tuanmu, B Parmentier, A Ranipeta, J Malczyk, and W Jetz. 2018. A suite of global, cross-scale topographic variables for environmental and biodiversity mapping. Scientific Data 5:180040. |
| Physical Characteristics | EarthEnvTopoMed_Elevation | Amatulli, G, S Domisch, M-N Tuanmu, B Parmentier, A Ranipeta, J Malczyk, and W Jetz. 2018. A suite of global, cross-scale topographic variables for environmental and biodiversity mapping. Scientific Data 5:180040. |
| Physical Characteristics | EarthEnvTopoMed_Slope | Amatulli, G, S Domisch, M-N Tuanmu, B Parmentier, A Ranipeta, J Malczyk, and W Jetz. 2018. A suite of global, cross-scale topographic variables for environmental and biodiversity mapping. Scientific Data 5:180040. |
| Physical Characteristics | EarthEnvTopoMed_TopoPositionIndex | Amatulli, G, S Domisch, M-N Tuanmu, B Parmentier, A Ranipeta, J Malczyk, and W Jetz. 2018. A suite of global, cross-scale topographic variables for environmental and biodiversity mapping. Scientific Data 5:180040. |
| Soil Characteristics | PelletierEtAl_  SoilAndSedimentary  DepositThicknesses | Pelletier, JD, PD Broxton, P Hazenberg, X Zeng, PA Troch, G-Y Niu, Z Williams, MA Brunke, and D Gochis. 2016. A gridded global data set of soil, intact regolith, and sedimentary deposit thicknesses for regional and global land surface modeling. Journal of Advances in Modeling Earth Systems 8:41-65. |
| Soil Characteristics | SG_Depth_to_bedrock | Hengl, T, J Mendes de Jesus, GBM Heuvelink, MR Gonzalez, M Kilibarda, A Blagotic, W Shangguan, MN Wright, X Geng, B Bauer-Marschallinger, MA Guevara, R Vargas, RA MacMillan, NH Batjes, JGB Leenaars, E Riberio, I Wheeler, S Mantel, and B Kempen. 2017. SoilGrids250m: Global gridded soil information based on machine learning. PLoS ONE 12:e0169748. |
| Soil Characteristics | SG_Sand_Content_005cm | Hengl, T, J Mendes de Jesus, GBM Heuvelink, MR Gonzalez, M Kilibarda, A Blagotic, W Shangguan, MN Wright, X Geng, B Bauer-Marschallinger, MA Guevara, R Vargas, RA MacMillan, NH Batjes, JGB Leenaars, E Riberio, I Wheeler, S Mantel, and B Kempen. 2017. SoilGrids250m: Global gridded soil information based on machine learning. PLoS ONE 12:e0169748. |
| Soil Characteristics | SG_SOC_Content_005cm | Hengl, T, J Mendes de Jesus, GBM Heuvelink, MR Gonzalez, M Kilibarda, A Blagotic, W Shangguan, MN Wright, X Geng, B Bauer-Marschallinger, MA Guevara, R Vargas, RA MacMillan, NH Batjes, JGB Leenaars, E Riberio, I Wheeler, S Mantel, and B Kempen. 2017. SoilGrids250m: Global gridded soil information based on machine learning. PLoS ONE 12:e0169748. |
| Soil Characteristics | SG_Soil_pH_H2O_005cm | Hengl, T, J Mendes de Jesus, GBM Heuvelink, MR Gonzalez, M Kilibarda, A Blagotic, W Shangguan, MN Wright, X Geng, B Bauer-Marschallinger, MA Guevara, R Vargas, RA MacMillan, NH Batjes, JGB Leenaars, E Riberio, I Wheeler, S Mantel, and B Kempen. 2017. SoilGrids250m: Global gridded soil information based on machine learning. PLoS ONE 12:e0169748. |
| Disturbance | ConsensusLandCover_Human_Development_Percentage | Tuanmu, M-N, and W Jetz. 2014. A global 1-km consensus land-cover product for biodiversity and ecosystem modelling. Global Ecology and Biogeography 23:1031-1045. |
| Disturbance | EsaCci_BurntAreasProbability | ESA Land Cover CCI project team; Defourny, P. 2016. ESA Land Cover Climate Change Initiative (Land_Cover_cci): Land Surface Seasonality Products. Centre for Environmental Data Analysis <https://catalogue.ceda.ac.uk/uuid/7c114fc6e2884c1f9ca107e7a502fdbf> |
| Disturbance | GHS_Population_Density | European Commission, Joint research centre, Friere, S, C Corbane, L Zanchetta, M Schiavina, P Politis, T Kemper, D Ehrlich, M Pesarest, L Maffenini, AJ Florczyk, M Melchiorri, and F Sabo. GHSL data package 2019: public release GHS P2019, Publications Office, 2019. https://data.europa.eu/doi/10.2760/290498. |

Table S1. Class, variable, and source for datasets used as independent niches axis for fungal occurrence data.

Table S2 (included as excel file). Current and future projected range sizes for all mycorrhizal fungal taxa as well as # of observations, niche hypervolumes, and % range shift.

| Mycorrhizal type | Taxonomic level | Variable | K | K  (p-value) | λ | λ  (p-value) |
| --- | --- | --- | --- | --- | --- | --- |
| EcM | Genus | Current range | **0.902** | **0.018** | **0.996** | **< 0.001** |
| EcM | Genus | Future range (ssp 126) | **0.903** | **0.017** | **0.998** | **< 0.001** |
| EcM | Genus | Future range (ssp 585) | **0.870** | **0.025** | **0.990** | **< 0.001** |
| EcM | Genus | % range shift (ssp 126) | 0.178 | 0.714 | 0.223 | 0.795 |
| EcM | Genus | % range shift (ssp 585) | 0.236 | 0.405 | 0.510 | 0.089 |
| EcM | Genus | Climatic niche hypervolume | **0.686** | **0.035** | **0.923** | **< 0.001** |
| AM | Genus | Current range | 0.214 | 0.933 | < 0.001 | 1.000 |
| AM | Genus | Future range (ssp 126) | 0.211 | 0.941 | < 0.001 | 1.000 |
| AM | Genus | Future range (ssp 585) | 0.206 | 0.950 | < 0.001 | 1.000 |
| AM | Genus | % range shift (ssp 126) | 0.271 | 0.796 | < 0.001 | 1.000 |
| AM | Genus | % range shift (ssp 585) | 0.218 | 0.904 | < 0.001 | 1.000 |
| AM | Genus | Climatic niche hypervolume | 0.593 | 0.222 | 0.389 | 0.795 |
| AM | Taxon | Current range | 0.109 | 0.079 | 0.170 | 0.312 |
| AM | Taxon | Future range (ssp 126) | 0.109 | 0.064 | 0.170 | 0.280 |
| AM | Taxon | Future range (ssp 585) | 0.110 | 0.055 | 0.162 | 0.274 |
| AM | Taxon | % range shift (ssp 126) | 0.079 | 0.769 | < 0.001 | 1.000 |
| AM | Taxon | % range shift (ssp 585) | 0.082 | 0.720 | < 0.001 | 1.000 |
| AM | Taxon | Climatic niche hypervolume | **0.123** | **0.026** | 0.351 | 0.066 |

Table S3. The phylogenetic signal of clematis niche hypervolume, current projected range, future

projected range, and modeled % range shifts for arbuscular mycorrhizal (AM) and

ectomycorrhizal (EcM) genera as well as AM fungal VTX. Blomberg’s K representing

branch-length divergence and Pagel’s λ representing clustering among closely related genera.

Statistical significance was determined via 1000 permutation and is indicated as : p<0.05(*),

p<0.01(**), and p<0.001(***).

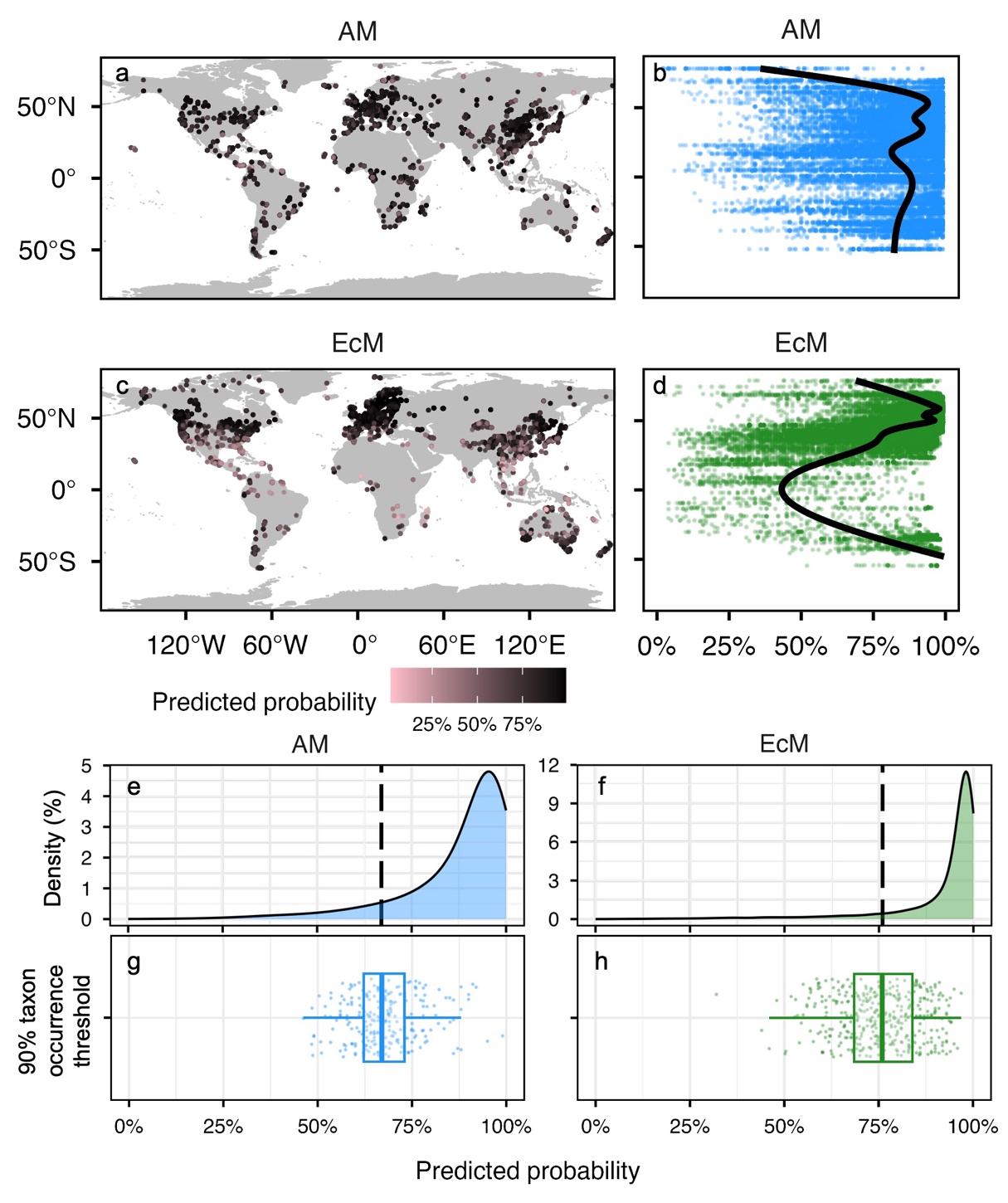

Figure S1. Modeled probability of occurrence for fungal taxa observations (a, c) with darker colors representing greater probability (model accuracy). Distribution of sample probabilities (b, d) for AM (blue) and EcM (green) fungal occurrences across latitude. Distribution of fungal occurrence threshold used to identify designate potentially suitable habitat from probability maps (e-h) with a dashed line representing the median probability threshold for assigning modeled occurrence for each mycorrhizal fungal guild. Thresholds were chosen using a decile approach where 90% of observed occurrences were captured in current maps at that probability.

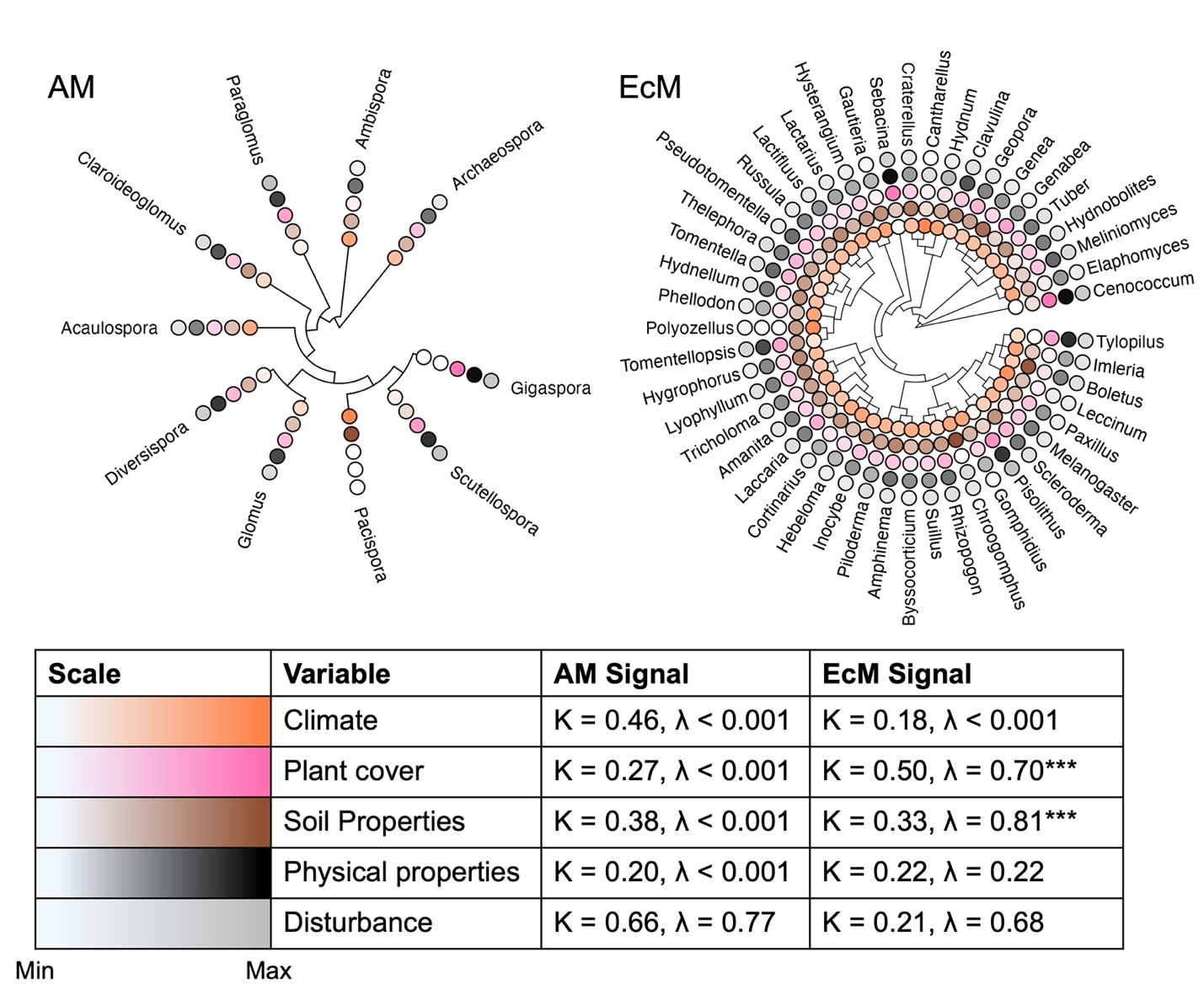

Figure S2. Phylogenetic trees for arbuscular mycorrhizal (AM, left) and ectomycorrhizal (EM,

right) genera. The relative feature importance of climate (orange), plant cover (pink), soil properties (brown), physical properties (black), and disturbance (grey) are shown in order at the

tips, represented by the color scales from white (minimum) to full color (maximum) for their

respective variables. The phylogenetic signal of these variables is shown in the bottom table,

with Blomberg’s K representing branch-length divergence and Pagel’s λ representing clustering

among closely related genera. Statistical significance was determined via permutation and is

indicated as : p<0.05(*), p<0.01(**), and p<0.001(***).

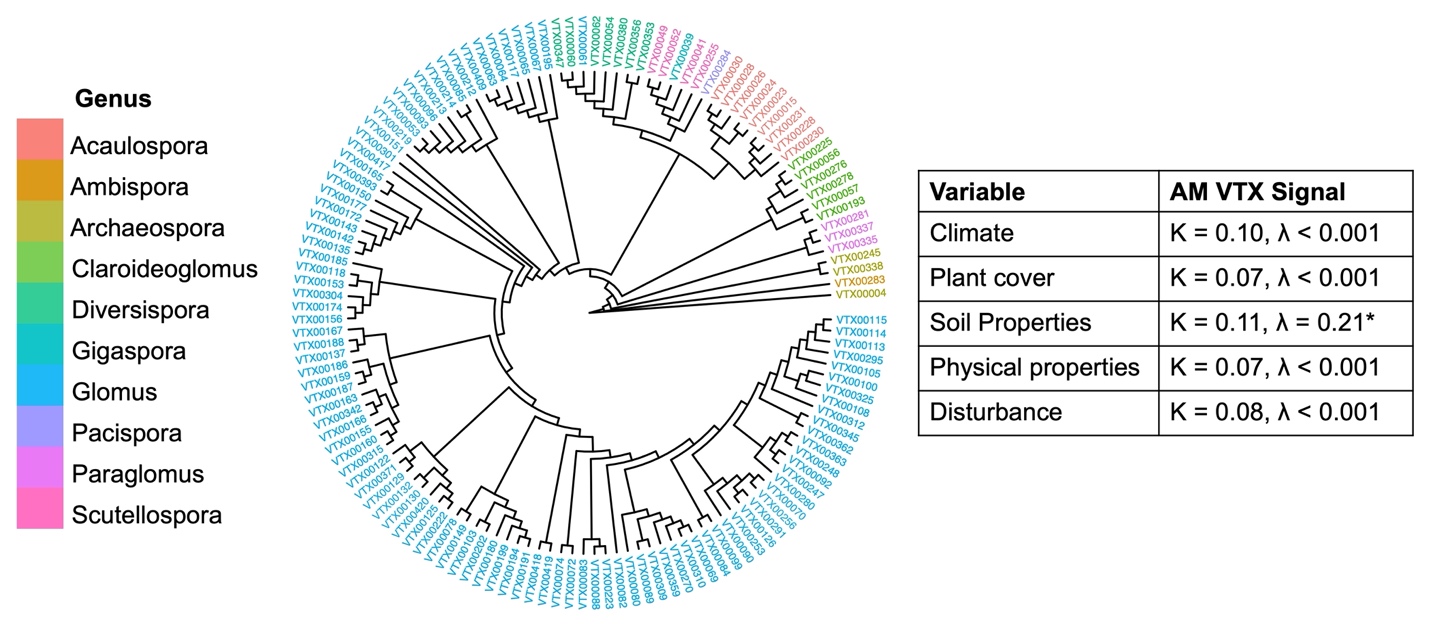

Figure S3. Phylogenetic tree of arbuscular mycorrhizal (AM) fungal VTX with tip labels colored

based on genus. The phylogenetic signal of relative feature importance variables are shown in

the right table, with Blomberg’s K representing branch-length divergence and Pagel’s λ

representing clustering among closely related genera. Statistical significance was determined via

permutation and is indicated as : p<0.05(*), p<0.01(**), and p<0.001(***).

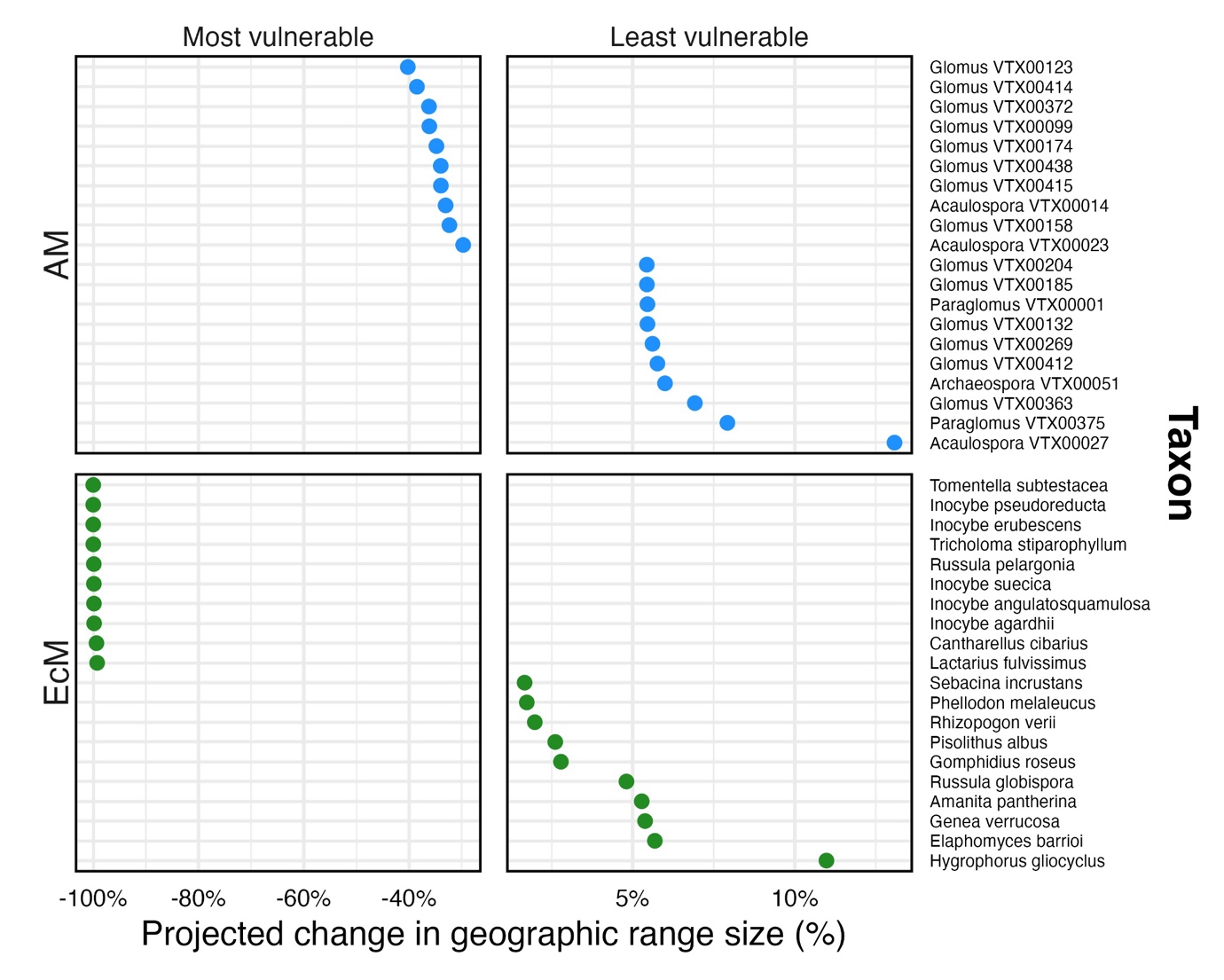

Figure S4. The 10 most and least vulnerable taxa from AM (blue) and EcM (green) fungal guilds based on projected change in geographic range size under the most sever climate scenario we used (ssp5-8.5).

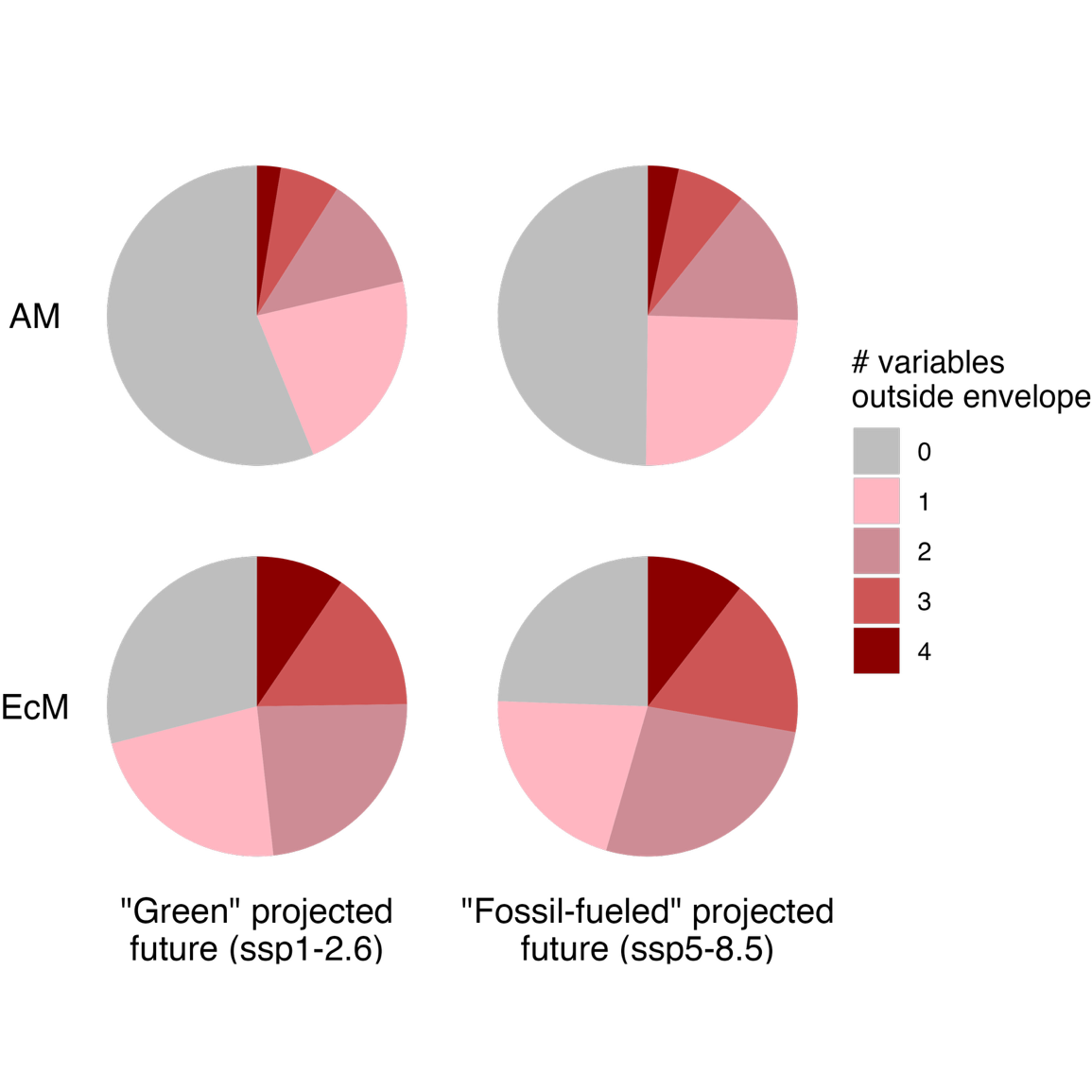

Figure S5. Pie charts showing the proportion of total areas and number of simulated future climate variables falling outside current climate envelopes for mycorrhizal taxa at the “best” case (ssp1-2.6) and “worst” case (ssp 5-8.5) climate scenarios.
